# A bacterial pathogen reveals a new mechanism of micron-scale nucleocytoplasmic transport

**DOI:** 10.64898/2026.11.25.769186

**Authors:** Hannah Kurka Margolis, Supratim Dey, George Minasov, Lauren E. Bird, Mohammad Sadik, Ingeborg M. Langohr, David Mankus, Jiayi He, Abigail K.R. Lytton-Jean, Oliver H. Voss, Karla J.F. Satchell, Rebecca L. Lamason

## Abstract

In addition to its canonical role as a selective barrier, the nuclear envelope is increasingly recognized as a dynamic structure, remodeled to integrate mechanical cues. Here, we discover that the bacterial pathogen *Rickettsia parkeri* deforms the nuclear envelope to an unprecedented degree to invade and exit human cell nuclei through large membrane protrusions. This extreme envelope remodeling is tolerated in part due to a bacterial protein that associates with the nuclear membrane and promotes nuclear infection, along with a mechanosensitive host enzyme that releases a signaling molecule from deformed nuclei. These findings establish a new mechanism of nucleocytoplasmic transport that redefines the size limits of nuclear envelope deformation cells can tolerate and resolves a long-standing question of how micron-scale cargo, such as bacteria, enter nuclei.

## Introduction

The nuclear envelope is a critical barrier that protects genetic material while simultaneously enabling communication and transport between the nucleus and cytoplasm. Although canonical transport is strictly regulated and size dependent, studying nucleocytoplasmic transport of pathogens has led to fundamental discoveries about the capacity of the nuclear envelope to regulate nuclear entry and egress. For instance, HIV transport through the nuclear pore complex (NPC) demonstrated unexpected NPC dilation that supports cargo up to ~60 nm in diameter (1), and herpes simplex virus (HSV) nuclear egress uncovered an NPC-independent transport process known as nuclear envelope budding (NEB) (2, 3) that was later shown to also enable ribonucleoprotein (RNP) complex egress from the nucleus (4, 5). While remarkable, these mechanisms are restricted to nanometer-scale cargo.

Despite being an order of magnitude larger than viruses, bacteria can also enter nuclei. Such intranuclear bacteria are typically considered quite rare, with most examples restricted to non-mammalian hosts such as amoebae (6), paramecia (7), and mollusks (8). A key exception to this is spotted fever group (SFG) *Rickettsia* spp., which have been observed in the nuclei of infected mammals (9), ticks (10–13), and tissue culture (14–18) for over 100 years. These tick-transmitted, neglected human pathogens can cause deadly vascular diseases and are becoming more common world-wide. Though largely considered cytosolic pathogens, prior reports suggested that nuclear infection by SFG *Rickettsia* spp. is frequent (9, 12) and that rickettsiae inside of the nuclei are motile (17, 18). Despite these observations, no known mechanism explains how large, micron-scale bacteria such as *Rickettsia* spp. cross the nuclear envelope.

As obligate, intracellular pathogens, SFG *Rickettsia* spp. must manipulate host cell machinery and membranes to ensure entry and dissemination through infected tissues (19). For example, surface-associated bacterial proteins (e.g., Sca2) polymerize host cell actin into tails that enable cytosolic motility (20). Prior work also showed that a secreted effector protein, Sca4, weakens intercellular tension to enable spread into neighboring cells (21). Thus, we reasoned that SFG rickettsiae may also actively manipulate the nuclear envelope to enter nuclei. Here, we present evidence that motile rickettsiae undergo a new mechanism of micron-scale nucleocytoplasmic transport that is promoted by a secreted bacterial protein.

## Results

### *Rickettsia parkeri* frequently enters nuclei of various human cell types where it undergoes actin-based motility

To understand how rickettsiae invade host nuclei, we first needed to determine the frequency of this phenomenon. We infected EA.hy926 (vascular endothelial-like cells, a physiologically relevant target) (14, 22, 23), A549, and U2OS cells (established infection models) (21, 24, 25) with the model SFG species *Rickettsia parkeri*. We then used concanavalin A and an antibody against the rickettsial surface protein OmpA to visualize the nuclear envelope and bacterial surface, respectively. We observed *R. parkeri* in the nuclei of all cell lines tested at 72 hours post infection (hpi). While the frequency of nuclear invasion varied by cell line from ~4% to over 20%, all frequencies were 10 to 100 times higher than those reported for the human bacterial pathogens *Burkholderia thailandensis* and *Shigella flexneri* (**Fig. 1A-B**) (26). By comparison, *Listeria monocytogenes*, another bacterial pathogen that undergoes actin-based-motility, was never observed in nuclei during infection (**fig. S1A-B**). Across the cell lines tested, the number of *R. parkeri* in each nucleus also varied from a single bacterium to as many as 50 to 100 bacteria densely packed within a single nucleus (**Fig. 1B**). Focused ion beam scanning electron microscopy (FIB-SEM) volumes further confirmed that bacteria are inside of, not adjacent to, nuclei and revealed *R. parkeri* with septa, suggesting that intranuclear *R. parkeri* can replicate (**Fig. 1C, movie S1**). We also noted that nuclear contents were absent where intranuclear bacteria were observed (**Fig. 1C**), suggesting that the bacteria exclude chromatin from their immediate vicinity. Yet *R. parkeri* infection was not associated with DNA damage (**fig. S2**).

**Fig 1.**
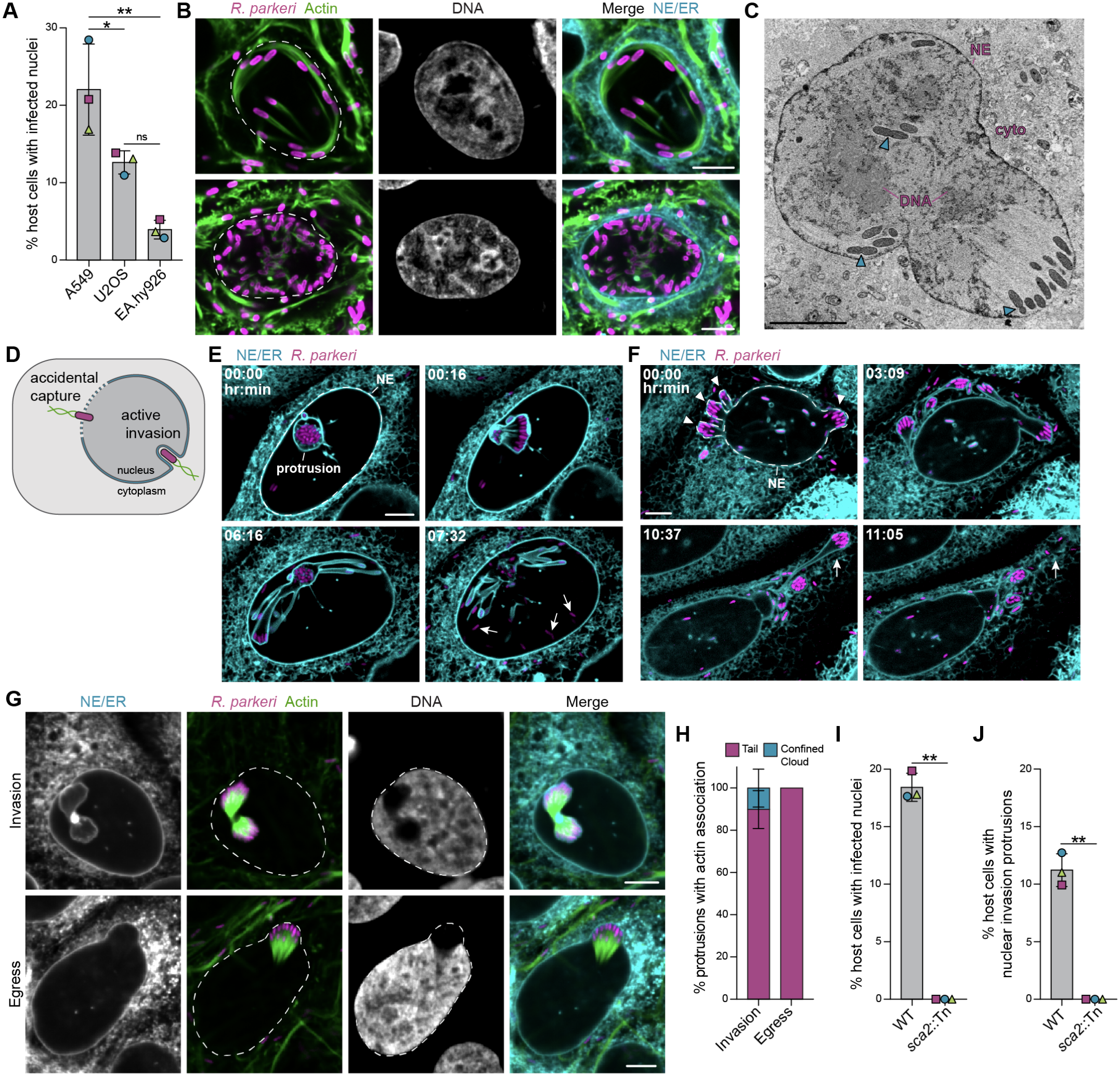
*R*. parkeri invade and exit human cell nuclei through membrane protrusions using actin-based-motility. (**A**) Frequency of *R. parkeri*-infected nuclei across human cell lines. Data are mean ± S.D. for n = 3 biological replicates (>90 infected cells/condition/replicate). Significance determined via one-way ANOVA with Dunnett’s multiple comparisons test. (**B**) Two representative images from (A) showing *R. parkeri* (magenta, anti-*Rickettsia*) in A549 cell nuclei (host DNA, grey, Hoechst), with F-actin (green, phalloidin), nuclear envelope and endoplasmic reticulum (cyan, concanavalin A). Dotted line, nuclear boundary. (**C**) Z slice from FIB-SEM volume of an *R. parkeri*-infected A549 cell with indicated cytoplasm (cyto), nuclear envelope (NE), dividing bacteria (cyan triangles), and displaced host genetic material (DNA). (**D**) Possible mechanisms of nuclear invasion. (**E**-**F**) Live-cell microscopy snapshots of mEmerald-Sec61 U2OS cells (cyan) infected with *R. parkeri* (BFP, magenta) invading (E) and egressing from (F) nuclei, with indicated NE (dotted line), protrusion (white triangles) and free *R. parkeri* in nucleoplasm or cytoplasm (white arrows). (**G**) Representative images from data in (H) showing *R. parkeri* invading and egressing from mEmerald-Sec61 U2OS nuclei (grey, Hoechst), with indicated F-actin tails (green, phalloidin), NE/ER (cyan, mEmerald-Sec61). Nuclear boundary, dotted line. (B-C, E-G) Scale bar, 5 µm. (**H**) Frequency of actin tails or clouds associated with *R. parkeri* in nuclear protrusions. Data are mean ± S.D for n = 3 biological replicates (>30 invasion, >15 egress events/replicate). (**I-J**) A549 cells infected with WT *R. parkeri* or *sca2*::Tn mutant, stained as in (A), and quantified for nuclear infection (I) or nuclear protrusions (J). Data are mean ± S.D. from n = 3 biological replicates (>100 infected cells/condition/replicate). Significance calculated via two-tailed Welch’s T-test. (A-B, G-J): samples harvested ~ 72 hpi. ^**^P ≤ 0.01, ^*^P ≤ 0.05, ns = not significant.

SFG *Rickettsia* spp. move throughout the host cell cytoplasm by polymerizing host cell actin into filaments at one of their cell poles (20). Using phalloidin to visualize F-actin, we observed intranuclear bacteria associated with long, straight actin tails that resembled those formed in the host cell cytoplasm (**Fig. 1B**). Time-lapse microscopy further confirmed that intranuclear *R. parkeri* were motile (**movie S2**), which further distinguishes them from nonmotile, intranuclear *B. thailandensis* (26). Together with the results above, these findings indicate that unlike other cytosolic bacterial pathogens, *R. parkeri* frequently invades nuclei and polymerizes actin tails to facilitate intranuclear movement.

### *R. parkeri* invade and exit human cell nuclei through membrane protrusions using actin-based motility

The nuclear envelope selectively transports nanometer-scale cargo while protecting genetic material. How micron-scale *R. parkeri* invades the nucleus was unclear. We considered two models: “accidental capture”, in which bacteria are passively trapped as the nuclear envelope reforms after host cell division (6, 26, 27), or “active invasion”, in which bacteria actively push into nuclei through protrusions at the nuclear envelope (**Fig. 1D**) (18). To distinguish between these models, we performed time-lapse microscopy of U2OS cells expressing mEmerald-Sec61 (to label the nuclear envelope and endoplasmic reticulum) (28) and infected with fluorophore-expressing *R. parkeri*. For all invasion events, we observed large *R. parkeri*-mediated protrusions at the nuclear envelope, with many spanning the diameter of the nucleus, before breaking to release the bacteria into the nucleoplasm (**Fig. 1E, movie S3**). To our surprise, bacteria also egressed from nuclei via large membrane protrusions that could extend to the cell periphery before subsequent release of the bacteria into the cytoplasm (**Fig. 1F, movie S4**). Both individual *R. par-keri* and groups of bacteria invaded and exited nuclei, with time to invade or egress ranging from ~20 minutes to over 24 hours (movie S5). Scanning transmission electron microscopy (STEM) revealed that *R. parkeri* push simultaneously on the inner and outer nuclear membranes when invading and egressing from host nuclei (fig. S3A-B). Bacteria were never observed between the inner and outer nuclear membranes, distinguishing *R. parkeri*-nucleocyto-plasmic transport from smaller-scale viral and RNP egress mechanisms, which rely on sequential deformation of single membranes (2–5, 29) (fig. S3C). These data also further distinguish *R. parkeri* from *B. thailandensis* and *S. flexneri*, which infrequently enter nuclei through a proposed accidental capture process (26). By contrast, we show that *R. parkeri* actively invades and exits nuclei via large membrane protrusions that are distinct from known forms of nucleocytoplasmic transport.

*R. parkeri*-induced nuclear protrusions were up to an order of magnitude larger than the 1 µm protrusions formed by these bacteria at the plasma membrane during cell-to-cell spread (21). As plasma membrane protrusions form without actin tails (21), we reasoned that deforming the more rigid nuclear envelope requires additional force from actin-based-motility. Indeed, over 80% of invading and all egressing bacteria within nuclear envelope protrusions had actin tails, and the remaining ~20% were out-lined by actin that appeared physically confined by the protrusion boundary (**Fig. 1G-H, fig. S4**). Additionally, both nuclear invasion protrusions and nuclear infection (i.e., bacteria free in the nucleoplasm) were abolished when cells were infected with a motility-deficient *R. parkeri* mutant (*sca2*::Tn) (30) (**Fig. 1I-J, fig. S5**). These data indicate that actin-based motility is essential for *R. parkeri* to form the large protrusions that drive nuclear invasion and egress, differentiating this process from cell-to-cell spread.

### A secreted bacterial protein localizes to nuclear protrusions and promotes *R. parkeri* nuclear invasion and egress

Because actin-based motility was necessary for *R. parkeri* to invade nuclei but insufficient for other motile cytosolic bacteria (fig. S1A), we hypothesized that proteins unique to *Rickettsia* spp. promote protrusion-mediated nuclear invasion. We focused on secreted effectors, which are proteins that pathogens translocate into the host cell to hijack various cellular processes. Using immunofluorescence microscopy, we screened the endogenous localization of several secreted effectors we recently discovered (31) and observed that the *Rickettsia*-specific secreted rick-ettsial factor E (SrfE) concentrated at the nuclear membrane of 85% and 95% of invasion and egress protrusions, respectively (**Fig. 2A, fig. S6A**). As expected, this pattern was lost in cells infected with an *R. parkeri* mutant strain lacking SrfE (*srfE*::Tn, **Fig. 2A-B, fig. S6A**) (32) and western blot confirmed SrfE was undetectable (fig. S6B).

**Fig 2.**
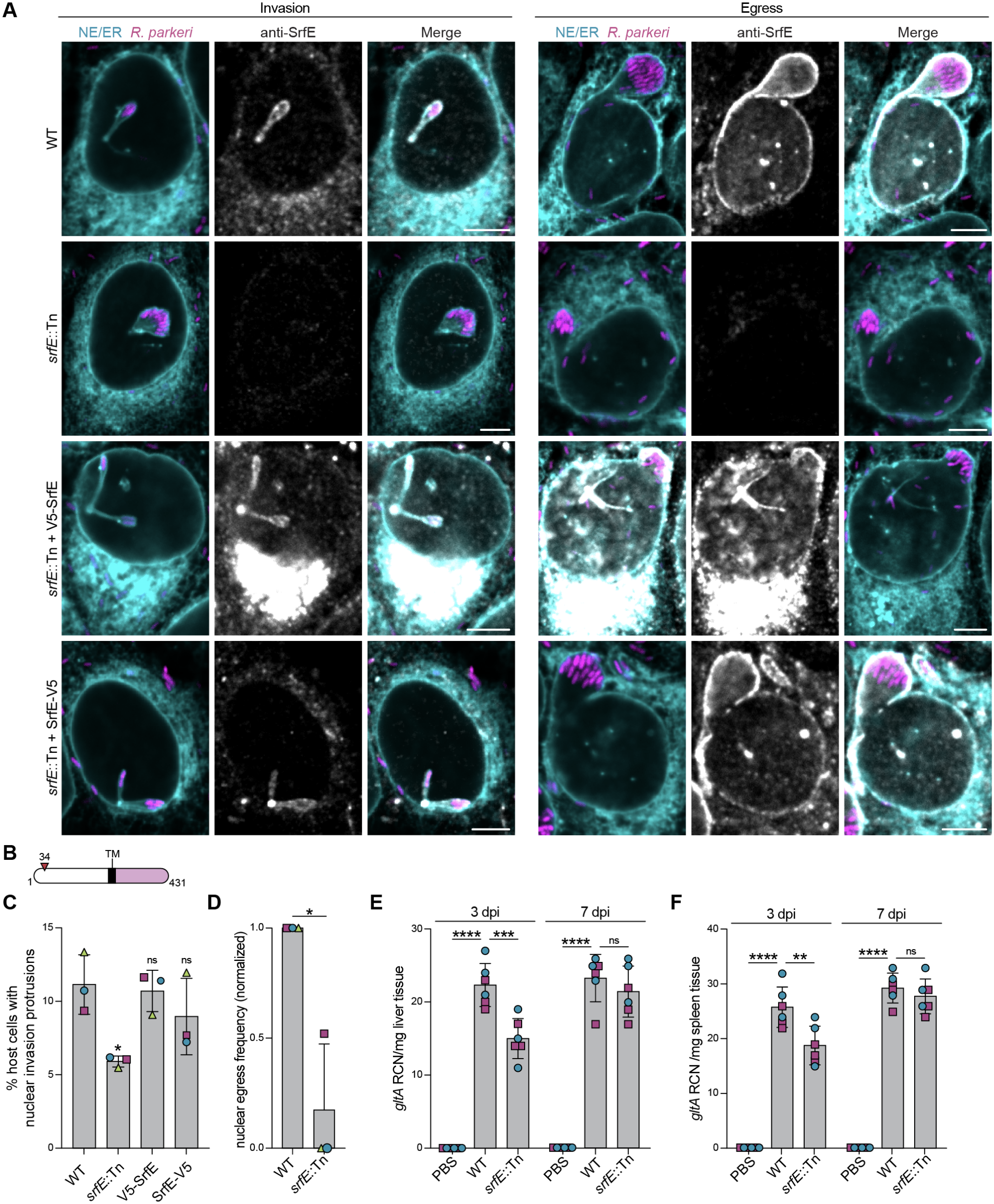
A secreted bacterial protein promotes *R. parkeri* nucleocytoplasmic transport and contributes to early stages of *in vivo* infection. (**A**) Representative images of SrfE localization to nuclear invasion and egress protrusions in mEmerald-Sec61 U2OS cells infected with WT, *srfE*::Tn, and complement *R. parkeri* strains (BFP, pseudocolored magenta) for ~72 h. NE/ER (cyan, concanavalin A); SrfE (grey, anti-SrfE); Scale bars, 5 µm. (**B**) SrfE protein schematic indicating N-terminal domain (white), transmembrane (TM) domain (black), C-terminal domain (light purple), and transposon insertion at indicated residue (red triangle). (**C**) Percent of infected A549 cells at ~72 hpi with nuclear invasion protrusions upon infection with indicated *R. parkeri* strains. Data are mean ± S.D. from n = 3 biological replicates (>300 infected cells/condition/replicate). Significance, relative to WT, calculated via one-way ANOVA with Dunnett’s multiple comparisons test. (**D**) Fraction of *R. parkeri* strains that egress infected mEmerald-Sec61 U2OS nuclei within a 12-hour window, normalized to WT for each replicate. Data are mean ± S.D. from n = 3 biological replicates (>50 infected cells/condition/replicate). Significance calculated via two-tailed Welch’s T-test. ns = not significant. (**E**-**F**) Bacterial burdens in livers (E) and spleens (F) of C57BL/6 mice injected intravenously with PBS or indicated *R. parkeri* strains, measured at indicated times by qPCR (relative copy number (RCN) of bacterial *gltA* normalized to host GAPDH and tissue weight). *gltA, R. parkeri* citrate synthase. Data are mean ± S.D. from n = 2 biological replicates (3 mice/condition/replicate). Significance calculated via one-way ANOVA with Dunnett’s multiple comparisons test, relative to WT. ^****^P ≤ 0.0001, ^***^P ≤ 0.001, ^**^P ≤ 0.01, ^*^P ≤ 0.05, ns = not significant.

SrfE has no sequence or structural homology outside the *Rickettsia* genus (31). However, it contains a predicted transmembrane (TM) domain immediately followed by positively charged residues, consistent with a Type I single-pass topology with the C-terminus exposed to the cytoplasm or nucleoplasm (Fig. 2B, fig. S7A-C). To test this we first complemented *R. parkeri srfE*::Tn with SrfE fused to the small epitope V5 tag at either terminus. Both constructs restored SrfE protrusion localization to varying degrees, though V5-SrfE also accumulated in the perinuclear region (**Fig. 2A, fig. S6A-B**). We then infected host cells with the complementation strains and stained for the V5 epitope after selectively permeabilizing with digitonin (permeabilizes the plasma membrane only) or Triton X-100 (permeabilizes all membranes). We found that only the C-terminal epitope remained accessible under digitonin permeabilization (fig. S7D-E). Together, these results indicate that secreted SrfE is a single pass transmembrane protein localized to nuclear invasion and egress protrusions with its C-terminal domain exposed to the host cell cytoplasm or nucleoplasm.

To test if SrfE regulates nuclear invasion or egress of *R. parkeri*, we infected A549 cells for 72 hours with wild-type, *srfE*::Tn mutant, or complemented *R. parkeri* strains and visualized the infections by immunofluorescence microscopy. We first noted that loss of SrfE did not significantly affect the frequency of host cells with infected nuclei (WT 15.15% ± 0.058; *srfE*::Tn 12.56% ± 3.40). We reasoned that these data could be explained by defects in both nuclear invasion and egress that would result in an unchanged steady state of nuclear infection frequency. Indeed, we observed a reduced frequency of cells with nuclear invasion protrusions upon *srfE*::Tn infection, and both complement strains restored this defect to near wild-type levels (**Fig. 2C**). Likewise, using time-lapse microscopy to capture rarer egress events, we found that the *srfE*::Tn mutant egressed from nuclei over 80% less frequently than wild-type *R. parkeri* (**Fig. 2D**). Taken together, our data suggest that the rickettsial secreted effector, SrfE, localizes to the membrane of nuclear protrusions and promotes both nuclear invasion and egress of *R. parkeri*.

### SrfE promotes early colonization *in vivo*

Having established a potential role for SrfE in tissue culture, we next wondered if SrfE contributes to infection *in vivo*. We therefore infected wild type C57BL/6 mice intravenously and quantified bacterial burden in spleen and liver at 3 and 7 days post infection (dpi) as previously done (33–35). We observed a modest (approximately 30%) but significant reduction in *R. parkeri srfE*::Tn in both the liver and spleen at 3 dpi, and this effect was gone by 7 dpi (**Fig. 2E-F**). Infection with the *srfE*::Tn strain also resulted in less splenomegaly at 3 dpi (fig. S8A-B) and reduced liver and spleen thrombosis compared to the wild-type strain (table S1). We noted mild hepatomegaly at 7 dpi, but not 3 dpi, with no significant difference between *R. parkeri* strains at either time point (fig. S8D-E). Histology revealed that although *R. parkeri* infection increased inflammation in the spleen and liver at both 3 and 7 dpi, there was no significant difference in amount of inflammation between strains in either tissue (fig. S8C, F). Taken together, these data suggest that, in addition to its role *in vitro*, SrfE contributes to early *R. parkeri* colonization *in vivo*.

### SrfE prevents host cell death associated with nuclear protrusion formation

To define the cellular basis of SrfE’s role in infection, and particularly in nuclear invasion, we returned to timelapse imaging of infected cells, this time aiming to quantify the outcome of nuclear protrusions. We observed that while only ~7.7% ± 5.5 of protrusions formed by wild-type *R. parkeri* led to host cell death within a 12-hour window of protrusion formation, loss of SrfE more than doubled the frequency of host cell death within the same window (**Fig. 3A-B**). Given the remarkable level of nuclear envelope deformation during *R. parkeri* nuclear invasion, we therefore wondered if SrfE suppresses host cell death induced by this process to promote nuclear invasion. In agreement with this hypothesis, we determined that significantly more host cells infected with the *R. parkeri srfE*::Tn mutant had activated caspase 3/7 activity than wild-type infected or uninfected cells, and that this phenotype could be complemented with SrfE-V5 (**Fig. 3C**). Moreover, treating infected host cells with the pancaspase inhibitor Q-VD-OPh rescued the nuclear invasion protrusion defect of the *srfE*::Tn mutant, while having no significant effect on the wild-type strain (**Fig. 3D**). Together, these data suggest that SrfE promotes nuclear invasion by preventing host cell death following nuclear invasion protrusion formation.

**Fig 3.**
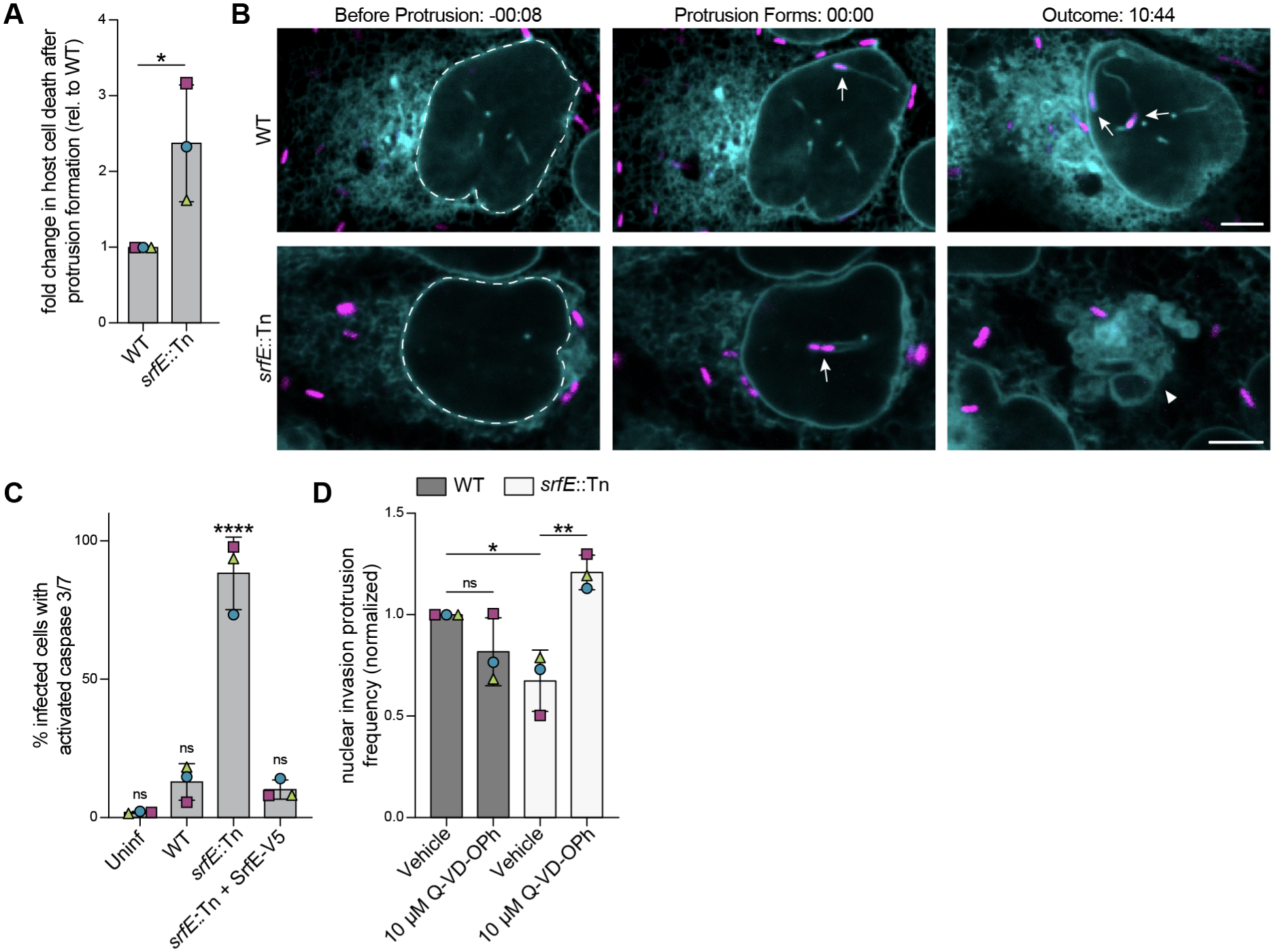
SrfE limits host cell death to promote nuclear invasion protrusions. (**A**)Fold change in host cell death within 12 hours of nuclear invasion protrusion formation by WT vs *srfE*::Tn *R. parkeri* quantified from live cell-microscopy. Data are mean ± S.D. from n = 3 biological replicates (>50 infected cells/condition/replicate). Significance calculated via Mann-Whitney test. (**B**) Live-cell microscopy snapshots of WT or *srfE:*:Tn *R. parkeri* (BFP, pseudocolored magenta) forming a persistent nuclear invasion protrusion (WT) or protrusion that ends in host cell death (*srfE*::Tn). Nuclear envelope (dotted white line); Nuclear protrusions (white arrows); dead host cell (white triangle); time in hour:min; scale bars, 5 µm. (**C**) Percent of infected A549 cells at ~72 hpi with activated caspase 3/7 signal. Data are mean ± S.D. from n = 3 biological replicates (>250 infected cells/condition/replicate). Significance calculated via one-way ANOVA with Dunnett’s multiple comparisons test, relative to WT. (**D**) Percent of infected A549 cells at ~72 hpi with nuclear invasion protrusions upon infection with indicated *R. parkeri* strains and treatment with Q-VD-OPh or vehicle control. Data are mean ± S.D. from n = 3 biological replicates (>250 infected cells/condition/replicate). Significance calculated via one-way ANOVA with Dunnett’s multiple comparisons test. ^****^P ≤ 0.0001, ^**^P ≤ 0.01, ^*^P ≤ 0.05, ns = not significant.

### The C-terminal domain of SrfE tetramerizes and binds a fatty acid

To complement our cellular assays, we sought to solve the structure of SrfE to understand the molecular basis of its function. Although we were unable to purify full-length SrfE or its N-terminal domain, we solved the structure of SrfE residues 277-403 (SrfE_CTD_) to 1.85 Å via x-ray crystallography (table S2). The structure revealed that the C-terminus of SrfE forms a tetramer, with each chain comprised of six helices which fold to form a paddle (**Fig. 4A**). Size exclusion chromatography coupled to multi angle light scattering (SEC-MALS) confirmed that SrfE_CTD_ exists in both a tetrameric and dimeric state in solution (fig. S9A-B). Neither the tetramer nor the monomers shared structural homology with known protein structures, precluding confident functional prediction.

**Fig 4.**
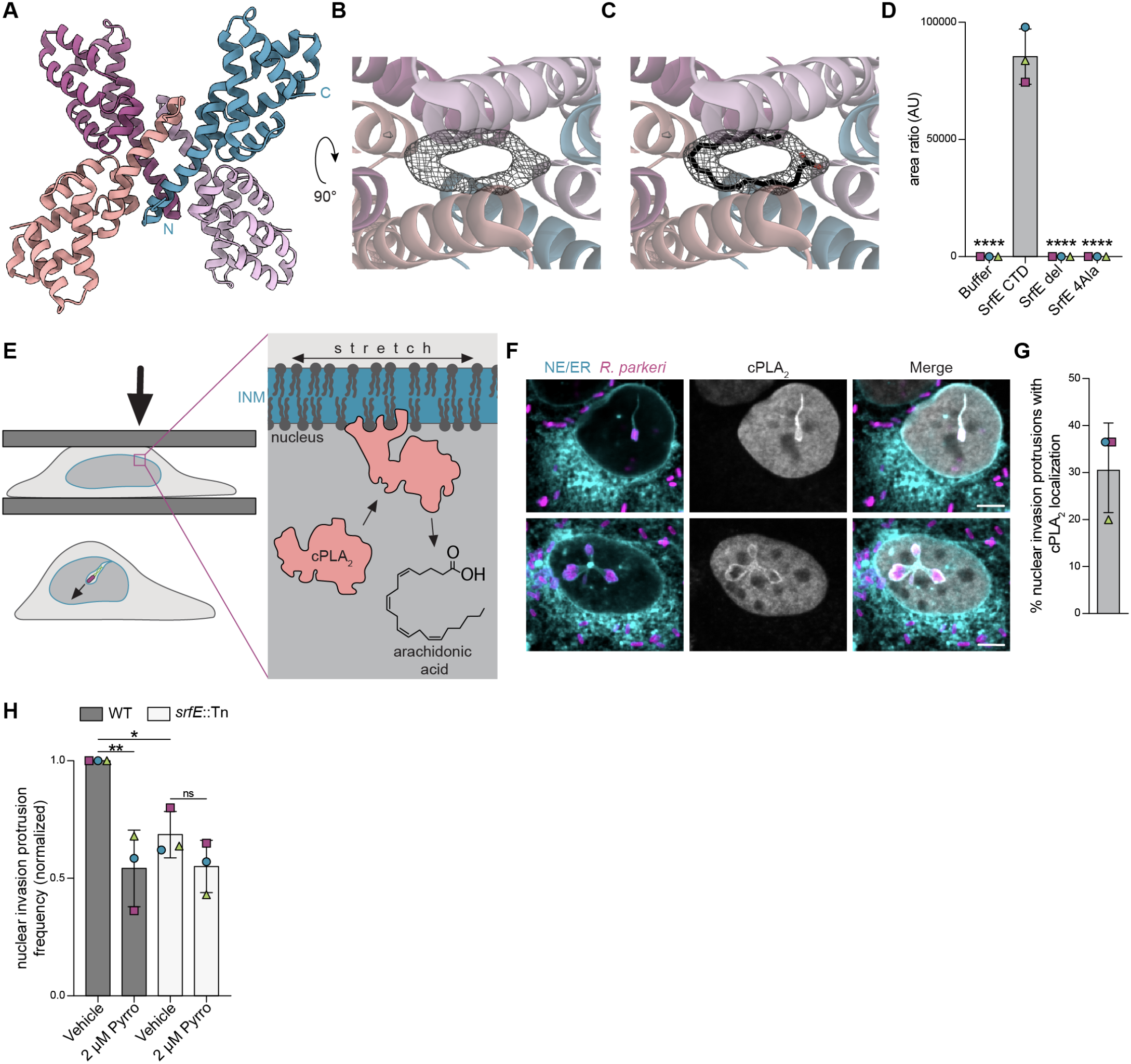
SrfE binds arachidonic acid and cPLA_**2**_ **activity promotes nuclear invasion protrusions**. (**A**)SrfE_CTD_ structure solved to 1.85 Å shown as a ribbon diagram. Each monomer colored separately, N and C termini indicated for blue monomer. (**B-C**) Density in tetramer interface (gray mesh) without (B) and with (C) a copy of arachidonic acid. Carbon in black, oxygen in red. (**D**) Copurification of arachidonic acid with indicated SrfE constructs from *E. coli* was quantified via mass spectrometry, using purification buffer as a negative control. Ratio is amount of arachidonic acid over amount of spiked in 13C1-palmitic acid loading control. Data are measured levels ± S.D. for n = 3 biological replicates. Significance, relative to WT was calculated via one-way ANOVA with Dunnett’s multiple comparisons test. (**E**) Model of arachidonic release via cPLA_2_ at sites of nuclear envelope tension during cellular compression or nuclear invasion. (**F-G**) Representative images (F) and quantification (G) of cPLA_2_ localization to nuclear invasion protrusions in mEmerald-Sec61 U2OS cells infected with WT *R. parkeri* strains (magenta, anti-*Rickettsia*) for ~72 h. NE/ER (cyan, concanavalin A); mKate2-cPLA_2_ (gray); Scale bars, 5 µm. Data in (G) are mean ± S.D. from n = 3 biological replicates (>50 invasion protrusions/condition/replicate). (**H**) Normalized percent of infected A549 cells at ~72 hpi with nuclear invasion protrusions upon infection with indicated *R. parkeri* strains and treatment with pyrrophenone or vehicle. Data are mean ± S.D. from n = 3 biological replicates (>250 infected cells/condition/replicate). Significance calculated via one-way ANOVA with Dunnett’s multiple comparisons test. ^****^P ≤ 0.0001, ^**^P ≤ 0.01, ^*^P ≤ 0.05, ns = not significant.

Intriguingly, the tetramer interface formed a hydrophobic pocket occupied by density that could not be assigned to the protein or any components of the crystallization buffer (**Fig. 4B**). To identify the source, we extracted crystals and subjected them to mass spectrometry analysis. This returned nine potential hits based on a cutoff of 4-log fold change over the crystallization buffer control, all of which were lipid in nature (table S3). However, only arachidonoyl-2-fluoroethylamide was a 20-carbon lipid, the length most consistent with the observed density. This molecule fit the density best with the 2-fluoro-ethyl group removed and when modeled in two possible confirmations, each with 0.5 occupancy (fig. S10A-B). As this lipid does not naturally occur in *E. coli* (from which we purified SrfE) nor was it a component of the crystallization buffer, we hypothesized that it copurified with SrfE out of the bacterial growth media.

Arachidononoyl-2-fluoroethylamide is closely related to arachidonic acid, an abundant polyunsaturated omega-6 fatty acid that is released from host cell membranes and converted into various signaling metabolites (36). Arachidonic acid fit within the observed density in our structure (**Fig. 4C, fig. S10C-D**) and copurified with SrfE_CTD_ as detected via mass spectrometry (Fig. 4D). Furthermore, deletion (SrfE_del_) or alanine substitution (SrfE_4Ala_) of the four residues that form the binding pocket at the tetramer interface (V285, L288, Y289, and F292) abolished this copurification (**Fig. 4D, fig. S10E**). These mutations were also sufficient to break the tetrameric interface of SrfE_CTD_ as determined by SEC-MALS (fig. S9A-D). Importantly, both mutant constructs were still properly folded based on melting curves (fig. S9E-J). Together, these data demonstrate that SrfE_CTD_ tetramerizes to bind arachidonic acid *in vitro*.

### cPLA_*2*_ activity promotes nuclear invasion protrusions in a SrfE-dependent manner

Physical deformation of the nuclear envelope stretches the phospholipid bilayer, recruiting the phospholipase cPLA_2_ to the inner nuclear membrane where it cleaves phospholipids at the sn-2 position to release arachidonic acid (**Fig. 4E**) (37–39). We hypothesized that the large nuclear envelope protrusions induced by *R. parkeri* would similarly recruit cPLA_2_, leading to release of arachidonic acid at the same sites occupied by SrfE (**Fig. 4E**). We stably expressed mKate2-cPLA_2_ (37) in mEmerald-Sec61 U2OS cells and observed that upon infection with *R. parkeri*, mKate2-cPLA_2_ was enriched at over 30% of nuclear invasion protrusions (**Fig. 4F-G**). No enrichment was observed elsewhere along the nuclear envelope, consistent with cPLA_2_ being specifically recruited to sites of nuclear invasion where it may locally release arachidonic acid.

Arachidonic acid plays signaling roles in various inflammatory pathways (36, 40) and may be connected to cell death (41). To test the importance of arachidonic acid binding to SrfE function, we attempted to complement the *R. parkeri srfE*::Tn strain with SrfE constructs carrying point mutations or deletions in the arachidonic acid binding site identified *in vitro*. However, mutating these residues severely impaired SrfE secretion from the bacterium (fig. S11). Therefore, we instead treated infected cells with pyrrophenone, a potent and specific inhibitor of cPLA_2_-mediated arachidonic acid release (42, 43). Pyrrophenone treatment reduced the frequency of nuclear invasion protrusions induced by wild-type *R. parkeri*, phenocopying the *srfE*::Tn mutant (**Fig. 4H**). Importantly, pyrrophenone did not significantly enhance the *srfE*::Tn mutant nuclear protrusion defect. These data suggest that arachidonic acid is locally released at nuclear protrusions by cPLA_2_ and supports nuclear invasion protrusions in a SrfE-dependent manner. Whether SrfE’s interaction with arachidonic acid is tied to its ability to prevent host cell death remains to be determined. Collectively, though, our data support a model in which *R. parkeri* actively invades the nucleus, using a combination of membrane-targeted host and bacterial proteins that support deformation of the nuclear envelope and passage of the bacterium into the nucleoplasm.

## Discussion

Fundamental discoveries of nucleocytoplasmic transport have been made by studying pathogen trafficking, such as NPC dilation to enable HIV nuclear entry (1) and nuclear envelope budding to enable HSV nuclear egress (2, 3). Here, we reveal how the bacterial pathogen *R. parkeri*, which is an order of magnitude larger than these viruses, can breach the nuclear envelope, uncovering a new mechanism of nucleocytoplasmic transport for micron-scale cargo. To cross the nuclear envelope through enormous membrane protrusions, *R. parkeri* manipulates multiple host cell processes, including actin polymerization (via Sca2), membrane deformation, and cell death (via SrfE). More broadly, nuclear invasion by *R. parkeri*, and the host cell manipulations required for its success, offer a rare opportunity to dissect how our cells respond to and regulate extensive nuclear envelope remodeling, chromatin displacement, and intranuclear actin, all of which are fundamental processes for gene regulation, development, and cell survival (45–47).

## Acknowledgments

This work was supported in part by the Koch Institute Support (core) Grant P30-CA14051 from the National Cancer Institute. We thank the Koch Institute’s Robert A. Swanson (1969) Biotechnology Center for technical support, specifically Peterson (1957) Nanotechnology for Integrative BioEM Core Facility (RRID:SCR_018674) This research used resources of the National Synchrotron Light Source II, a U.S. Department of Energy (DOE) Office of Science User Facility operated for the DOE Office of Science by Brookhaven National Laboratory under Contract No. DE-SC0012704. Use of LS-CAT resources for data collection supported by the Michigan Economic Development Corporation and the Michigan Technology Tri-Corridor (Grant 085P1000817). Access to LS-CAT and computational resources is supported by the Northwestern Structural Biology Facility, which is funded in part by the Robert H. Lurie Comprehensive Cancer Center grant NCI P30CA060553. This research also used the Mass Spectrometry Core in Research Resources Center of University of Illinois at Chicago. We thank the Metabolomics Core at UMass Chan Medical School (RRID:SCR_027036) for assistance with experimental design, metabolite extraction, LCMS analysis, data processing and interpretation. We wish to acknowledge use of instrumentation in the Biophysical Instrumentation Facility (BIF) at MIT. We are grateful to Ryan Miller, Alexandra Wilson, David Bartel, Roberto Vazquez Nunez, Seychelle Vos, Dezmond Cole, Erin Goley, Ted Hackstadt, Daniel Portnoy, Chris Paddock, Noah Taylor, Ben Velez, Iain Cheeseman, Chris Obara, and Jennifer Lippincott-Schwartz for reagents. We thank Roberto Vazquez Nunez and Seychelle Vos for assistance with protein purification. The content is solely the responsibility of the authors and does not necessarily represent the official views of the National Institutes of Health. This manuscript is the result of funding in whole or in part by the National Institutes of Health (NIH). It is subject to the NIH Public Access Policy. Through acceptance of this federal funding, NIH has been given a right to make this manuscript publicly available in PubMed Central upon the Official Date of Publication, as defined by NIH.

## Author contributions

Conceptualization: HKM, RL Formal Analysis: HKM, GM Funding Acquisition: HKM, AKRL, OHV, KJFS, RL Investigation: HKM, SD, GM, LB, IML, DM, JH, MS, OHV Methodology: HKM, IML, DM, MS, AKRL, OHV Project Administration: AKRL Supervision: HKM, OHV, KJFS, RL Visualization: HKM, KJFS, RL Writing – original draft: HKM, RL Writing – review & editing: HKM, SD, GM, LB, IML, DM, MS, OHV, KJFS, RL

## Data, code, and materials availability

Structure and structural statistics have been deposited to the Protein Data Bank (www.rcsb.org) with accession code 35YA. The ligand identification and arachidonic acid binding mass spectrometry data generated in this study have been deposited in the public proteomics repository MassIVE (www.massive.ucsd.edu) under accession codes MSV000104114 and MSV000104150, respectively. All other data are available in the manuscript or the supplementary materials.

## Supplementary Materials

Figs. S1 to S11

Tables S1 to S3

Movies S1 to S5

## Methods

### Human cell line propagation and generation

A549 human lung epithelial, U2OS bone osteosarcoma, and Vero kidney epithelial cell lines were obtained from the University of California, Berkeley Cell Culture Facility (Berkeley, CA). EA.hy926 endothelial and U2OS mEmerald-Sec61beta cells were gifts from Daniel Swale (University of Florida) and Chris Obara/Jennifer Lippincott-Schwartz (Janelia Research Campus), respectively. All human cell lines were maintained in Dulbecco’s modified Eagle’s medium (DMEM; Gibco 11965118) supplemented with 10% fetal bovine serum (FBS). Vero cells were maintained in DMEM supplemented with 5% FBS. Cell lines were confirmed to be negative for mycoplasma by the Koch Institute Preclinical Modeling facility using MycoAlertTM PLUS Assay (Lonza LT07-710).

U2OS mEmerald-Sec61beta cPLA_2_-mKate2 cells were made by transfecting U2OS mEmerald-Sec61beta cells with cPLA_2_-mKate2 (Addgene 163844) and sleeping beauty transposase (kindly provided by the Bartel Lab, White-head Institute). Cells were selected using 2 µg/mL puromycin.

### Bacteria strain generation and propagation

All *R. parkeri* strains were made from the parental Portsmouth strain (kindly provided by Chris Paddock, Centers for Disease Control) and propagated for tissue culture infections in Vero cells grown in DMEM supplemented with 2% FBS at 33°C, as previously described (21). The *srfE*::Tn *R. parkeri* mutant was kindly provided by Thomas Burke (University of California Irvine). To make strains, *R. parkeri* was transformed with pRAM18d plasmid variations (see below) via electroporation and clonally isolated by plaquing on Vero cell monolayers overlayed with agarose, as previously described (48). Stocks were titered via plaque assay and stored in aliquots at −80°C. Spectinomycin (50 µg/mL) or rifampin (200 ng/mL) were included in media to select for transformants and to ensure plasmids were maintained in experiments.

*R. parkeri* strains were partially purified and used for intravenous mouse infections as described previously (35, 49).

*L. monocytogenes* strain 10403S expressing GFP (kindly provided by Daniel Portnoy, University of California Berkeley) were grown in brain heart infusion (BHI) broth at 30°C.

### Plasmid construction

Constructs for SrfE complement strains were made by amplifying nt 1113891 - 1115897 (SrfE with flanking intergenic regions, SrfE: MC1_06530; GenBank AFC75335.1; RefSeq MC1_RS05980 / WP_014411051.1) from wild-type *R. parkeri* strain Portsmouth (accession: NC_017044) and restriction cloned into puc19 (HKM_D03Fw and Rv) with SacI and SalI. NEB-uilder HiFi DNA Assembly Master Mix (New England BioLabs E2621) was used to add the N-(HKM_D06Fw and Rv) and C-(HKM_D07Fw and Rv) terminal V5 tag with linker. The resulting constructs were amplified out of pUC19 (HKM_D03Fw and Rv) and restriction cloned with SacI and SacII into a pRAM plasmid providing BFP expression and spec resistance (kindly provided by Erin Goley).

Constructs for mutated SrfE complement strains were made by mutagenizing the above C-terminal V5-tagged construct in pUC19 with Q5 Hot Start High Fidelity 2X Master Mix (New England BioLabs M0494) (HKM_D16Fw and Rv; HKM_D17Fw and Rv). Resulting constructs were amplified and restriction cloned into the same pRAM plasmid as above.

The SrfE_CTD_ construct for purification was made by codon optimizing the sequence of *srfE* for *E. coli*, synthesizing DNA, and cloning into onboarded vector pMCSG9 (50) (p6xHis-MBP-SrfE-C) by Twist Bioscience (South San Fransisco, CA). Mutagenesis of the p6xHis-MBP-SrfE-C plasmid was performed with Q5 Hot Start High Fidelity 2x Master Mix and primer pairs HKM_D09Fw and Rv; HKM_D11Fw and Rv.

Primers used for cloning:

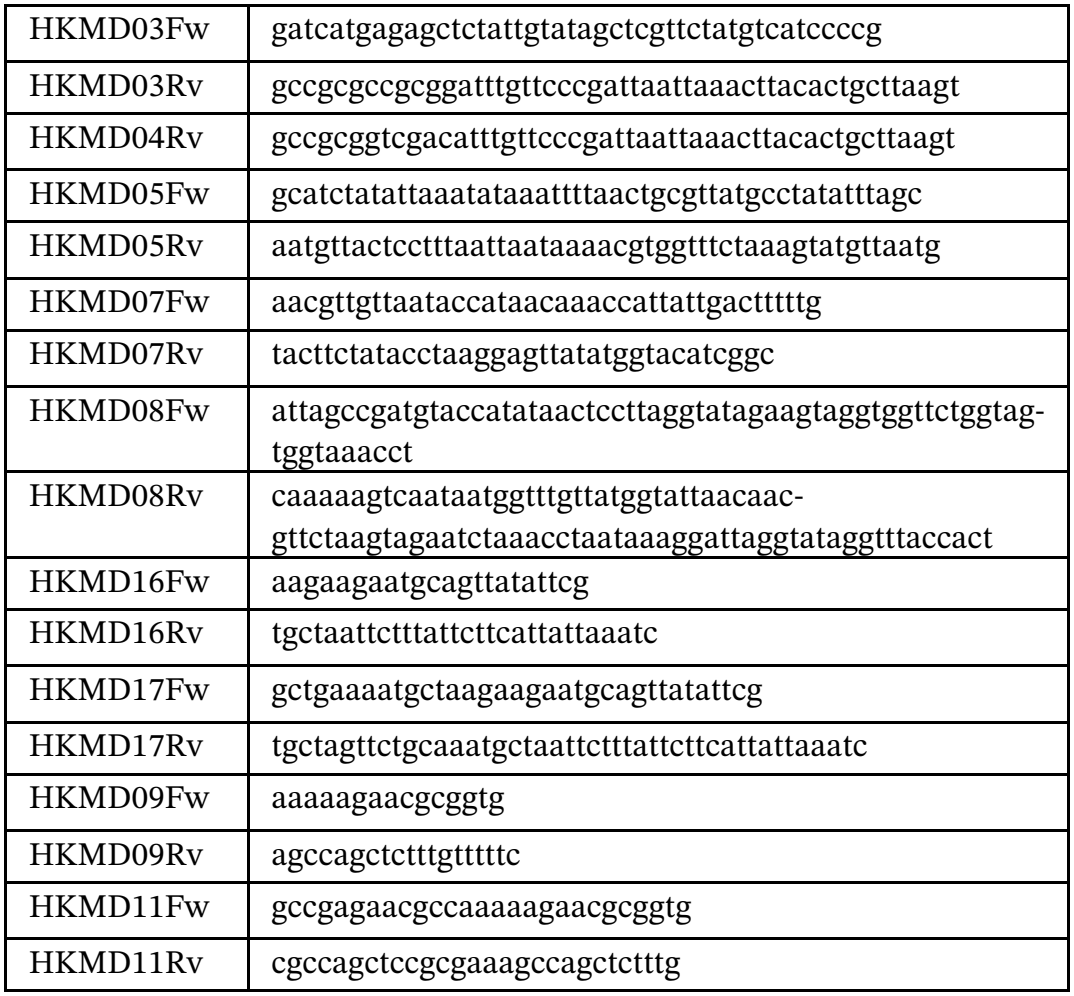

### Infections for confocal immunofluorescence microscopy and analysis

All immunofluorescence microscopy was performed as follows, with variations for individual experiments indicated below. Indicated host cells were seeded on glass #1.5 coverslips (Fisherbrand 12-541-001) in 24-well tissue culture treated plates (Genesee Scientific 25-107) and incubated for at least 16 hours at 37°C before infection. To infect, *R. parkeri* in BHI or *L. monocytogenes* in PBS were added to seeded cells and spinfected at 200 x g for 5 min at RT. Infections were performed such that all wells received an equal volume of BHI or PBS. Infections were incubated at 33°C. To maintain ER morphology, infected cells were fixed without exposure to air by removing half of the media from each well, mixing it 1:1 with 16% paraformaldehyde (PFA, Ted Pella 18505), and adding this mixture back to each well for 4% final PFA concentration. Cells were fixed 15 minutes (if no endogenous fluorophores) to 1 hour (if endogenous fluorophores were present in bacteria or cells) protected from light at RT and washed 2 times with 1X PBS. Samples were treated and stained at RT as follows: quenched with 0.1 mM glycine for 10 min, washed 3 times with 1X PBS, permeabilized with 0.5% Triton-X 100 in PBS for 5 min, washed 3 times with 1X PBS, blocked for at least 30 min with 2% BSA/PBS, stained with primary antibodies diluted in block solution for 1 hour, soaked 3 times for 5 min with 1X PBS, stained with secondary antibodies and cell stains diluted in block solution for 1 hour, soaked 3 times for 5 min with 1X PBS, and mounted using ProLong Gold antifade (Invitrogen P36930).

Images were acquired on an Olympus IXplore Spin microscope with Yokogawa CSU-W1 spinning disk and an ORCA-Flash4.0 sCMOS camera using either 60x UPlanSApo (1.30NA) or 100x UPlanSApo (1.35 NA) objectives. All image analysis was performed in ImageJ.

#### Nuclear infection and invasion protrusion assays

Cells were seeded at 2×10^5^ to 3×10^5^ cells/mL and infected with *R. parkeri* or *L. monocytogenes* at indicated MOIs for 72 hours. Images were burden matched for analysis by max projecting each image, determining the percent image area occupied by bacteria (CellProfiler) and approximate cell density (hand count). Images were counted by hand in ImageJ as follows: nuclei were first counted using only the Concanavalin A channel, *R. parkeri*, actin, and Concanvalin A channels were used to count the number of counted nuclei that had bacteria free in the nucleus and/or fully wrapped in a nuclear invasion protrusion.

##### Cell types (Fig. 1A-B)

Cells were infected at an MOI of 0.023-0.035 with wild-type *R. parkeri* for 72 hours and stained with Hoechst (Invitrogen 113570, 1:1K), anti-OmpA (14-13, kindly provided by Ted Hackstadt, 1:400), Phalloidin-AlexaFluor555 (Invitrogen A30106, 1:400), and Concanavalin A Conjugate AlexaFluor647 (Invitrogen C2142, 1:400)

##### Wild-type vs sca2::Tn (Fig. 1I-J, fig. S5)

Cells were infected at an MOI of 0.023-0.05 with wild-type *R. parkeri* expressing GFP or *R. parkeri* strain *sca2*::Tn for 72 hours in media with 200 ng/mL rifampin and stained as for Fig1A-B (Cell Types).

##### R. parkeri vs L. monocytogenes (fig. S1)

Cells were infected with wild-type *R. parkeri* at an MOI of 0.023 for 72 hours or *L. monocytogenes* 10403S strain expressing GFP at an MOI of 1-1.5 for 5 hours and stained as for Fig1A-B (Cell Types).

##### Wild-type, srfE::Tn, and complement strains (Fig. 2C, 3D, 4H)

Cells were infected at an MOI of 0.0025 with *R. parkeri* strains expressing BFP (wild-type, *srfE*::Tn, *srfE*::Tn + V5-SrfE, *srfE*::Tn + SrfE-V5) for 72 hours in media with 50 µg/mL spectinomycin and stained with Phalloidin-AlexaFluor647 (Invitrogen A22287, 1:400) and Concanavalin A Conjugate AlexaFluor488 (Invitrogen C11252, 1:100). For caspase inhibition (Fig 3D), cells were pre-treated for at least 1 hour at 37°C with 50 µg/mL spectinomycin and 10 µM pancaspase inhibitor Q-VD-OPh (APExBio A1901) or DMSO vehicle control before bacteria were directly added for spinfection. For cPLA_2_ inhibition assay (Fig 4H), cells were pre-treated for at least 1 hour at 37°C with 50 µg/mL spectinomycin and 2 µM pyrrophenone (Sigma-Aldrich 5305380001) or vehicle control before bacteria were directly added for spinfection. Media was changed approximately every 24 hours to maintain fresh pyrrophenone.

#### Actin tail frequency (Fig. 1G-H, fig. S4)

U2OS cells stably expressing mEmeraldSec61-beta were infected at an MOI of 0.017 with *R. parkeri* expressing BFP for 72 hours in media with 200 ng/mL rifampin and stained with Concanavalin A Conjugate AlexaFluor488 (1:100), Phalloidin-AlexaFluor555 (1:100), and Sytox Deep Red (Invitrogen S11380, 1:2K). Images were counted by hand in ImageJ as follows: the *R. parkeri* and Concanavalin A channels were used to identify bacteria in nuclear invasion and egress protrusions before using the actin channel to identify tails (at least one bacterial length from the bacterial pole), confined protrusions (actin adjacent to bacteria less than a bacterial length), or no tail (no actin adjacent to bacteria).

#### yH2ax (fig. S2)

A549 cells were infected at an MOI of 0.017 with *R. parkeri* expressing GFP for 72 hours in media with 200 ng/mL rifampin and stained with anti-yH2ax (Sigma 05-636, 1:400) and Concanavalin A Conjugate AlexaFluor647 (1:400). Nuclei were traced in ImageJ using the Concanavalin A channel, cells were z-projected, and the mean intensity of yH2ax was measured.

#### Secreted SrfE staining (Fig. 2A, fig. S6A)

U2OS cells stably expressing mEmeraldSec61-beta were infected at an MOI of 0.01 with *R. parkeri* strains expressing BFP (wild-type, *srfE*::Tn, *srfE*::Tn + V5-SrfE, *srfE*::Tn + SrfE-V5) for 72 hours in media with 50 µg/mL spectinomycin and stained with anti-SrfE polyclonal antibody (1:400) (31) for 3 hours at 37°C and Concanavalin A Conjugate AlexaFluor488 (1:100). Block solution was 10% normal goat serum (Jackson ImmunoResearch 005-000-121) in 2% BSA/PBS. Images were counted by hand in ImageJ as follows: the *R. parkeri* and Concanavalin A channels were used to identify bacteria in nuclear invasion and egress protrusions before using the SrfE channel to identify protrusions with SrfE signal.

#### SrfE orientation (fig. S7D-E)

U2OS cells stably expressing mEmeraldSec61-beta were infected at an MOI of 0.01 with *R. parkeri* strains expressing BFP (wild-type, *srfE*::Tn + V5-SrfE, *srfE*::Tn + SrfE-V5) for 72 hours in media with 50 µg/mL spectinomycin. Samples were permeabilized with either 0.5% Triton-X 100 in 1X PBS for 5 min at RT or 30 µg/mL digitonin in 1X PBS for 3 min at RT. For epitope-tagged SrfE detection, samples were stained with anti-V5 (Invitrogen 45-0705, 1:200) for 3 hours at 37°C. For controls, samples were stained with anti-calnexin (Abcam ab22595, 1:200) or anti-Lamin A/C (Sigma MABT1341, 1:400) for 3 hours at 37°C. Block solution was 10% normal goat serum in 2% BSA/PBS. Images were burden matched as described above and z projected. Cell boundaries were traced in ImageJ (using the Concanavalin A channel) and the mean intensity was measured for the channel of interest. The mean digitonin and triton values upon infection with wild-type *R. parkeri* (no V5 tag) were subtracted from the values upon infection V5-tagged SrfE strains to account for nonspecific staining before the ratio was taken.

#### Activated caspase 3/7 activity (Fig. 3C)

A549 cells were infected at an MOI of 0.0025 with *R. parkeri* strains expressing BFP (wild-type, *srfE*::Tn, *srfE*::Tn + V5-SrfE, *srfE*::Tn + SrfE-V5) for 72 hours in media with 50 µg/mL spectinomycin. After 4 hours, media was replaced to contain 50 µg/mL spectinomycin and 5 µM BioTracker NucView 488 Green Caspase Dye (Sigma-Aldrich SCT101). Samples were stained with Concanavalin A Conjugate AlexaFluor647 (1:400). Images were burden matched as described above. Images were counted by hand in ImageJ as follows: the caspase channel was set to remove nonspecific cytoplasmic background signal and minimize signal in uninfected samples, the Concanavalin A and *R. parkeri* channels were used to count the number of infected cells, and the caspase dye channel was used to identify cells with signal in the nucleus (counted positive).

#### cPLA_2_ localization to nuclear invasion protrusions (Fig. 4F-G)

U2OS cells stably expressing mEmeraldSec61-beta and cPLA_2_-mKate2 were infected at an MOI of 0.017-0.025 with *R*.*parkeri* expressing BFP for 72 hours in media with 200 ng/mL rifampicin and stained with Concanavalin A Conjugate AlexaFluor488 (1:100). Images were counted by hand in ImageJ as follows: the *R. parkeri* and Concanavalin A channels were used to identify bacteria in nuclear invasion protrusions and the cPLA_2_ channel was used to identify protrusions with signal. Cells were counted as positive if cPLA_2_ signal at protrusions was at least 1.5x greater than the surrounding nucleoplasm signal (within one bacterial length of the protrusion).

### Electron microscopy

For focused ion beam scanning electron microscopy (FIB-SEM) volume (Fig. 1C, movie S1), A549s were seeded at 2.5×10^5^ cells/mL as described for confocal immunofluorescence microscopy experiments and infected with *R. parkeri* expressing GFP at an MOI of 0.092 in media containing 50 µg/mL spectinomycin for 48 hours at 33°C. Cells were washed 1X with PBS and fixed with 4% PFA, 0.2% glutaraldehyde in 1X PBS for 1 hour protected from light. Staining was done as described above using Hoechst (1:1000). Samples were screened on a Zeiss LSM 880 confocal to locate regions of interest. Regions of interest were mapped in both fluorescent and brightfield contrast to assist in correlation. Coverslips were triple washed with 0.1M sodium cacodylate buffer, incubated in 1% osmium tetroxide and 1.25% potassium ferrocyanide in 0.1M sodium cacodylate for 1 hour at 4°C, washed 4 times with ultrapure water, incubated with aqueous 1% thiocarbohydrazine for 20 minutes at RT, washed 4 times with ultrapure water, incubated in 1% osmium tetroxide for 30 minutes, washed 3 times in 50mM maleate buffer (pH 5.15), and incubated overnight in 2% uranyl acetate in 50mM maleate buffer at RT in the dark. Samples were dehydrated through a graded series of ethanol, (30, 50, 70, 85, 90, 95, 100% 3 times), transitioned into acetone (50/50, 70/30, 100% 2 times), and infiltrated with Durcupan resin (30/70, 50/50, 70/30, 100% 2 times). Polymerized blocks were removed from coverslips using liquid nitrogen immersion to thermally contract the plastic and glass, and pressed blocks from the BEEM capsules using a purpose-built capsule press. Block faces were imaged with a Leica S9i stereomicroscope to correlate with the earlier fluorescent and brightfield imaging. Block faces were coated with approximately 10nm of platinum-palladium alloy using a Denton BenchTop Turbo evaporator and mounted into a Zeiss 540 Cross-beam FIB-SEM. The block face was imaged using the SEM beam and correlated to light microscopy data using the Zeiss ATLAS 5 software. The identified cell was imaged with a 10nm pixel size and a 10nm slice thickness for a volume of approximately 25µm by 24µm by 12µm depth.

For scanning transmission electron microscopy (STEM) imaging (fig. S3A-B), U2OS mEmerald-Sec61beta cells were seeded at 3 to 5×10^5^ cells/mL as described for confocal immunofluorescence microscopy experiments and infected with BFP-expressing *R. parkeri* at an MOI of 0.002-0.02 in media containing 200 ng/mL rifampin for 72 hours at 33°C. Cells were fixed without exposure to air as described for “confocal immunofluorescence microscopy” using a final concentration of 4% PFA, 0.2% glutaraldehyde in 1X PBS for 1 hour protected from light. Coverslips were washed 2 times with 1X PBS and screened on a Zeiss LSM 880 confocal to locate regions of interest. Coverslips were tripled washed with 0.1M sodium cacodylate buffer, incubated in 1% osmium tetroxide and 1.25% potassium ferrocyanide in 0.1M sodium cacodylate for 1 hour at 4°C, rinsed 2 times with 0.1M sodium cacodylate, rinsed 2 times with 50mM maleate buffer (pH 5.15), and incubated overnight in 2% uranyl acetate in 50mM maleate buffer at RT in the dark. Samples were then en bloc stained in uranyl acetate, dehydrated, infiltrated, and resin embedded as described above and mounted to inverted BEEM capsules filled with resin placed strategically around the regions of interest. Blocks were polymerized for at least 48 hours in a 60°C oven and the coverslips were removed using liquid nitrogen as described above. Blocks were pressed from the BEEM capsules using a purpose-built capsule press, the block face was manually trimmed to a trapezoidal shape approximately 1.5mm in length per side using a razor blade, and 60 nm ultrathin sections were cut using a DiATOME diamond knife on a Leica UC7 ultramicrotome. Short ribbons were collected on formvar-carbon copper slot grids and imaged on a Zeiss 540 Crossbeam FIB-SEM with inserted STEM detector at 30kV and 500pA.

### Time-lapse confocal microscopy

All time-lapse microscopy was performed with variations in the following protocol as indicated for individual experiments below. U2OS mEmerald-Sec61beta cells were seeded at 2.5×10^5^ to 5.5×10^5^ cells/mL in Nunc Lab Tech II Chambered Coverglass dishes (ThermoFisher Scientific 155409) pre-treated with 20 µg/mL human fibronectin (Millipore Sigma FC010) for 1 hour at RT and incubated for at least 16 hours at 37°C before infection. Cells were infected in media containing antibiotic as indicated below by spinfection at 200 x g for 5 min at RT. Infections were incubated at 33°C. Before imaging, wells were washed twice with phosphate buffered saline (PBS) and 200 µL of FluoroBrite DMEM (Thermo Fisher Scientific A1896701) supplemented with 10% FBS and containing appropriate antibiotic was added. Movies were acquired on an Olympus Ixplore Spin microscope system using an environmental chamber set to 5% CO2 and 33°C. Images were taken every 7 or 8 minutes. Movies were bleach corrected using the histogram matching feature within Image J and annotated using Adobe Premier Pro.

For Fig. 1E-F and movies S2-5, cells were infected at an MOI of 0.1-0.9 with *R. parkeri* expressing BFP in media with 200 ng/mL rifampin for at least 24 hours at 33°C before imaging. For Fig. 3A-B, cells were infected at an MOI of 0.2-2 with *R. parkeri* strains expressing BFP (wild-type, *srfE*::Tn) in media with 50 µg/mL spectinomycin for at least 2 hours at 33°C before imaging.

### Western blotting

A549s were seeded at 8×10^5^ cells/mL in 24-well tissue culture treated plates and incubated for at least 16 hours at 37°C before infection. Cells were spin-fected at 200 x g for 5 min at RT at an MOI of 0.012 with *R. parkeri* strains expressing BFP (wild-type, *srfE*::Tn, *srfE*::Tn + V5-SrfE, *srfE*::Tn + SrfE-V5) in media with 50 µg/mL spectinomycin for 72 hours at 33°C. Infected wells were gently washed 1X with PBS, trypsinized for 5 min at 33°C, and centrifuged at 2400 x g for 5 min at RT. Pellets were resuspended in selective lysis buffer (50 mM HEPES pH 7.9, 150 mM NaCl, 10% v/v glycerol, 1% v/v IGE-PAL supplemented with protease inhibitors (Sigma-Aldrich P1860), incubated on ice for 20 min, and centrifuged at 11,300 x g for 10 min at 4°C. Pellets were washed 2 times without resuspension with 1X PBS and resuspended in loading dye (50 mM Tris-HCl pH 6.8, 2% SDS, 10% glycerol, 0.1% bromophenol blue, 5% β-mercaptoethanol). Supernatant was passed through a 0.22-µm cellulose acetate filter (Thermo Fisher Scientific F2517-1) by centrifugation at 6700 x g for 10 min at 4°C and denatured with loading dye. Lysates were boiled and detected via Western blot using mouse anti-V5 (Invitrogen 46-0705; 1:2000), rabbit anti-SrfE polyclonal antibody (1:2000), and mouse anti-RpoB (BioLegend 663905; 1:2000).

### Animal infections

For mouse infections, male, wild-type C57BL/6 mice were exposed to a heat lamp for about 5 min prior to being moved into a mouse restrainer. The tail was sterilized with 70% ethanol, and 200 µL of either rickettsial suspension (10^7^ pfu in PBS) or PBS alone was administrated into the lateral tail vain as described previously (35, 49). Animals were monitored for any signs of discomfort and euthanized using CO_2_ followed by cervical dislocation at days 3 and 7 post-infection for specimen collection. Livers and spleens were collected and weighed.

### Burden analysis

qPCR was performed on RNA isolated from spleen and liver as previously described (35, 49, 51, 52). Briefly, organs were homogenized in Trizol reagent (Invitrogen 15-596-018) via bead beating, RNA was extracted using the Quick-RNA miniprep kit (ZymoResearch R1055), and cDNA was synthesized from 200 ng using the GoScript™ Reverse Transcription System (Promega A5001) according to the manufacturer’s instructions. qPCR amplification and detection were performed on a QuantStudio 3 Real-Time PCR System (Applied Biosystems by Thermo Fisher) using iQ™ SYBR^®^ Green Supermix (BioRad 170-8882), 2 µL cDNA (5 ng/ µL) and 1 µM each of the following oligonucleotides for rickettsial (housekeeping) citrate synthase gene (*gltA*) and host (housekeeping) *GAPDH* gene (see primers below). Cycling conditions were as follows: 1 cycle at 95°C for 3 min; 40 cycles at 95°C for 15 sec, 55°C for 15 sec, and 72°C for 20 sec; and 1 cycle to generate the dissociation curve. Melting curve analyses were performed at the end of each run to ensure that only one product was amplified. The relative copy number (RCN) of *gltA* expression was normalized by the expression of *GAPDH* and amount of organ tissue (≤ 10 mg spleen, ≤ 50 mg liver) as described previously (35, 49).

Primers used for PCR:

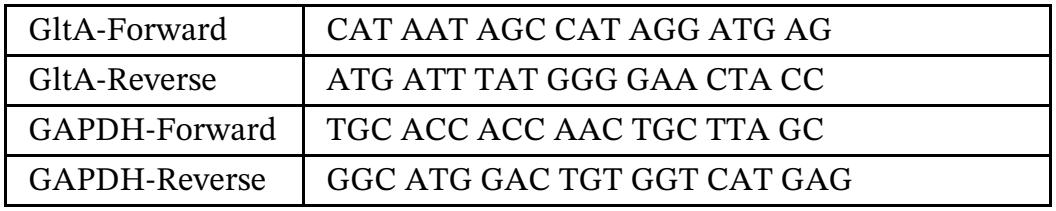

### Histology processing and analysis

Mouse organs were fixed in 10% neutral buffered formalin for 24 hours and transferred to 70% ethanol. Histology staining was performed by HistoWiz Inc using their standard operating procedure. Samples were processed, embedded in paraffin, and sectioned at 5 µm thickness Hematoxylin and eosin (H&E)-stained sections of liver and spleen were evaluated by light microscopy by a board-certified veterinary pathologist. Two histopathologic parameters were assessed in each tissue. Inflammation was scored semi-quantitatively on a 5-point ordinal scale: 0 = absent, 1 = minimal, 2 = mild, 3 = moderate, and 4 = marked. Grades were assigned based on the severity and extent of the inflammatory infiltrate, ranging from rare inflammatory cells (minimal) to involvement of <25%, 25-50%, or >50% of the tissue (mild, moderate, and marked, respectively). Thrombosis was recorded as a binary variable and scored as present (+) or absent (-) for each tissue and animal.

### Protein purification

For biochemical analysis and arachidonic acid detection, constructs were expressed and purified as previously described (31).

For structural determination, the 6xHis-MBP-TEV-SrfE (277-403) construct was expressed in *E. coli* BL21(DE3) (Magic) cells as previously described (*53*), grown in High Yield M9 SeMet media (Medicilon Inc. MD045004) supplemented with 200 µg/mL ampicillin and 50 µg/mL kanamycin, and induced overnight with 0.25 mM IPTG at 25°C. Pelleted cells were resuspended in lysis buffer (25 mM Tris pH 8.3, 0.5 M NaCl, 10% v/v glycerol, 0.1% IGEPAL CA-630, 1 mM TCEP) and sonicated at 50% amplitude, in 5 s (on) × 10 s (off) cycles for 20 min on ice. The lysate was cleared by centrifugation at 18,000 × g for 45 min at 4 °C, the supernatant was collected, and the protein was loaded onto Ni-NTA resin pre-equilibrated with 5 column volumes (CV) of loading buffer (25mM Tris-HCl pH 8.3, 500 mM NaCl, 1 mM TCEP, 10 mM imidazole and 10% glycerol), washed with 10 CV of wash buffer (25mM Tris-HCl pH 8.3, 500 mM NaCl, 1 mM TCEP, 30 mM imidazole and 10% glycerol), and eluted with elution buffer (25mM Tris-HCl pH 8.3, 500 mM NaCl, 1 mM TCEP, 300 mM imidazole and 10% glycerol). Protein was dialyzed overnight at 4°C (25 mM Tris-HCl pH 8.3, 500 mM NaCl, 1 mM TCEP, 10 mM imidazole and 10% glycerol) with TEV protease to remove the 6xHis-MBP tag. Cleaved protein was further purified by a second round of Ni-NTA affinity chromatography and then size exclusion chromatography using Superdex 200 16/600 column (10mM Tris HCl pH 8.3, 250mM NaCl, 1mM TCEP and 2% glycerol).

### Crystallization and structure determination

SrfE_CTD_ was concentrated to 9.4 mg/ml in buffer A and screened for crystallization as 2 µL sitting drops (1 µL protein: 1 µL reservoir solution) in 96-well SWISSCI (High Wycomb, England) UVXPO MRC 2-Lens Crystallisation Plates using commercial Classics II, PEG’s II and AmSO4 (QIAGEN 130723, 130716, 130705) crystallization screens by vapor diffusion at 19°C. Most screen conditions from AmSO4 screen yielded crystals. Diffraction quality crystal of the protein used for the final data collection grew from the condition with 0.1 M citric acid (pH 5.0), 1.6 M ammonium sulfate (AmSO4 screen, #E8) and was flash cooled in liquid nitrogen for data collection using 2 M LiSO4 as cryoprotectant.

Data sets were collected at beamline 17-ID-1 of the National Synchrotron Light Source II (NSLS-II) at Brookhaven National in collaboration with the Life Sciences-Collaborative Access Team (LS-CAT) at the Advanced Photon Source, Argonne National Laboratory. Images were indexed, integrated, and scaled using HKL-3000–2.3.15 (54). Data quality and structure refinement statistics appear in Table SX The structure was determined by Molecular Replacement Method implementing AlphaFold3 (55) model (model ID AF-Q9ZCI0-F1) using Phaser (56). Initial model went through several rounds of refinement in REFMAC-5.8.0431 (57) and manual model corrections in Coot (0.9.4.1) (58). The water molecules were generated using ARP/wARP −8.0 (59) followed by additional rounds of refinement in REF-MAC-5.8.0431. The ligand was modeled as an arachidonoyl amine corresponding to N-(2-fluro-ethyl) arachidonoyl amine from mass spectrometry results. The quality control of the models during refinement and for the final validation of the structures was done using MolProbity (60). Structures were deposited to the RCSB PDB (https://www.rcsb.org/) with the assigned PDB code 35YA. Structure figures were prepared with UCSF ChimeraX-1.12 and Pymol-3.1.3.1.

### Ligand identification

Protein crystals from the same well used for x-ray data collection (approx. 5) were pooled, washed twice in buffer (10 mM Tris HCl pH 8.3, 250 mM NaCl, 1 mM TCEP and 2% glycerol), dissolved in 10 mM ammonium bicarbonate, denatured at 98°C for 10 min, passed through a 5 kDa cut off concentrating column, and sent for mass spectrometry analysis to the Mass Spectrometry Core facility at University of Illinois Chicago. The above buffer and crystallization screen condition (0.1 M citric acid pH 5.0, 1.6 M ammonium sulfate) were also sent as controls.

Ice-cold methanol and chloroform (1:2 v/v) were added to each sample followed by 5 min of sonication and 30 min on ice with occasional vortexing. Ice-cold water was added at a final ratio of 8:4:3 chloroform:methanol:water and samples were incubated on ice for 10 min, then centrifuged at 10k RPM for 5 min at 4°C. The aqueous (compound extracts) and organic layer (lipid extracts) were extracted for separate analysis as follows.

MS analysis of the lipid extracts (lower phase, organic layer) was performed using an Agilent 6545 Q-TOF LC–MS system controlled by the Agilent Mass Hunter acquisition software. The mass spectrometer was operated in 2 GHz extended dynamic range mode employing precursor ion analysis for relative quantification experiments in positive ion mode. Internal references were used for calibration. Mobile phase A was 90:10 H2 O:methanol v/v with 10 mm ammonium acetate plus 0.5 mm ammonium fluoride and B was (5:3:2) isopropanol:methanol: acetonitrile v/v/v with 10 mm ammonium acetate plus 0.5 mm ammonium fluoride. The resuspended lipid extracts were loaded onto a 2.1 × 100 mm Agilent Poroshell C18, 2.7 µm column (Agilent Technologies Inc., Santa Clara, CA, USA) and separation performed using an Agilent 1290 UPLC system using the following gradient: 70% B at 0–1 min, 86% B at 3.5–10 min, 100% B at 11–17 at a flow rate of 300 µL/ min. A post column equilibration time of 5 min was used for all runs. Source parameters were as follows: gas temp (200 °C), drying gas (11 L/min), nebulizer (35 psi), sheath gas temp (350 °C), sheath gas flow (12 L/min), VCap (3000 V), and fragmentor (145 V). Data were collected for relative quantification using a scan speed of 4 MS spectra per second. Pooled samples were used in lipid database creation, and an iterative MS/MS workflow was performed in the Mass Hunter acquisition software across multiple injections of the pooled samples with a scan speed of 10 MS and 3 MS/MS spectra per second.

MS analysis of the compound extracts (upper phase, aqueous layer) was performed using an Agilent 6545 Q-TOF and 1290UPLC system controlled by the Agilent Mass Hunter acquisition software. The mass spectrometer was operated in 2 GHz extended dynamic range mode employing precursor ion analysis for relative quantification experiments in positive/negative ion modes. Internal references were used for calibration. Mobile phase A was 0.1% formic acid in H2O and B was 0.1% formic acid in ACN. The samples were loaded onto a 2.1 × 100 mm Agilent Poroshell C18, 1.9 µm column (Agilent Technologies Inc., Santa Clara, CA, USA) and separation performed using an Agilent 1290 UPLC system at a flow rate of 300 µL/ min. A post column equilibration time of 4 min was used for all runs. Source parameters were as follows: gas temp (300 °C), drying gas (11 L/min), nebulizer (35 psi), sheath gas temp (350 °C), sheath gas flow (12 L/min), VCap (3000 V), and fragmentor (145 V). Data were collected for relative quantification using a scan speed of 3 MS spectra per second.

Raw LC–MS/MS data were used to create a fragmentation-based (MS/MS) library containing m/z precursors and retention times for all MS/MS identified lipids (see figure 2 and 3) using the Lipid Annotator software (Agilent Technologies Inc., Santa Clara, CA, USA) and the following settings: lipid species for positive ([M + H]+, [M + Na]+, [M + NH4]+, [M + H − H2 O]+, [M + Na − H2 O]+, [M + NH4 − H2 O]+), Q-Score ≥ 60, and mass deviation ≤ 10 ppm. Raw LC–MS data files were processed using the Profinder software (vB.10.00, Agilent Technologies Inc., Santa Clara, CA, USA) with the library based targeted extraction. Here, molecular features were extracted for peaks ≥ 5000 counts in the positive mode ([M + H]+, [M + Na]+, [M + NH4]+, [M + H − H2 O]+, [M + Na − H2 O]+, [M + NH4 − H2 O]+) using an isotope model of common organic molecules. The resulting compound list was further filtered for those having an absolute height of ≥10,000 counts, quality score of ≥60, and compounds having two or more isotopes present. Additionally, retention time for each compound was aligned to ±0.1 min using a mass accuracy window of ≤5.0 ppm and peaks integrated using the Agile integrator in the Profinder software (level 2 identification). Each of the integrated peaks was manually reviewed for retention time and fragmentation matching. The processed data file was then exported and imported into the Mass Profiler Professional software (v15.1, Agilent Technologies Inc., Santa Clara, CA, USA) where each data set was analyzed separately. All compound abundance values were baseline corrected to the median abundance.

Raw LC–MS data files were processed using the Profinder software (vB.10.00, Agilent echnologies Inc., Santa Clara, CA, USA). Here, molecular features were extracted for peaks ≥ 3000 counts in the positive mode (5000 counts for negative mode) using an isotope model of common organic molecules. The resulting compound list was further filtered for those having an absolute height of ≥5000/7000 counts, quality score of ≥60, and compounds having two or more isotopes present. Additionally, retention time for each compound was aligned to ±0.1 min using a mass accuracy window of ≤5.0 ppm and peaks integrated using the Agile integrator in the Profinder software. The annotation of the compounds was conducted against METLIN Compound database. If multiple hits were found for a feature, the compound candidate with the highest score is chosen for the annotation of the feature.

### Detection of arachidonic acid with purified SrfE_CTD_

SrfE_CTD_, SrfE_del_, and SrfE_4Ala_ were purified as described above and sent to University of Massachusetts Chan Medical School Metabolomics Core. Water (Thermo Fisher Scientific W64), isopropanol (Fisher Chemical A461), methanol (Fisher Chemical A456), and acetonitrile (Fisher Chemical A955) were all OPTIMA LCMS grade. Formic acid was LCMS grade (Thermo Scientific Fisher Chemical A117). Chloroform was HPLC grade (Fisher Chemical C606SK). Protein preps were thawed on ice and 20 µL (approx. 3mg/ml) was combined with 100 µL 10mM ammonium bicarbonate. Samples were denatured at 65°C for 10 minutes, combined with 1 mL of cold 2:1 chloroform:methanol (with 0.5µM 13C1-Palmitic acid as internal standard) and sonicated for 5 minutes. Samples were incubated on ice for 30 minutes with occasional vortexing, combined with 180 µL cold water, incubated on ice for 10 minutes with occasional vortexing, and spun down at max speed in a tabletop centrifuge at 4°C for 10 minutes. The lower organic phase was then removed and dried down in a refrigerated speed vac for storage at −80°C until resuspension.

Dried samples were reconstituted in 1:1 acetonitrile:isopropanol and loaded into individual LC-MS vials. Fatty acids were examined using a Vanquish liquid chromatography (LC) platform coupled to a Orbitrap IQ-X Tribrid mass spectrometer equipped with a heated electrospray ionization (HESI) probe (Thermo Electron). LC separation was performed with an Acquity Premier HSS T3 column (1.8µm, 100mm X 2.1mm; Waters Corporation) using a gradient of solvent A (10 mM ammonium acetate in 60:40 acetonitrile:water) and solvent B (10 mM ammonium acetate in 88:10:2 isopropanol: acetonitrile: water). Flow rate was 300 µL/min. The LC gradient was: 0 min, 10% B; 2 min, 10% B; 16 min, 60% B; 21 min, 100% B; 22 min, 100% B; 24 min, 10% B. Column temperature was set to 50°C. The autosampler temperature was 15°C, and the injection volume was 3 µL. Source settings were 2300V for negative mode; sheath gas was set to 40 Arb, Aux gas 8 Arb, sweep gas 1 Arb and the ion transfer tube and vaporizer temperature were set to 300°C and 225°C, respectively. Mass spectra were acquired in negative ionization mode in parallel reaction monitoring for 303.233 m/z (arachidonic acid) and 256.2363 m/z (13C1-palmitic acid). For the MS1 acquisition, the resolution was set to 60,000, RF lens was 50%, the automatic gain control (AGC) target was 2×105, the maximum injection time was 100 msec, and the scan range was m/z 70-900 m/z. For MS/MS, the ions in each full scan were isolated in the quadrupole with a 1.5m/z window, fragmented with a normalized HCD stepped collision energy of 15, 30, 45, 75, 100 units, and analyzed at a resolution of 30,000 with an AGC target of 5×104 and a maximum injection time of 54 msec. External calibration was performed every 7 days.

Peaks were integrated in TraceFinder (Thermo) using a 5 ppm mass tolerance and matched to retention times determined with chemical standards. Area ratios were generated by dividing the peak area of arachidonic acid by the peak area of the internal standard 13C1-palmitic acid.

### Size exclusion chromatography-multi angle light scattering (SEC-MALS)

SrfE_CTD_, SrfE_del_, and SrfE_4Ala_ were dialyzed into new buffer (10 mM Tris pH 7.5, 250 mM NaCl, 1 mM TCEP), diluted to 0.5 mg/mL, and spun down at 14,100 x g for 15 minutes at 4°C. Samples were run on a WTC-030S5 size exclusion chromatography analytical column. Elution profiles and molar masses were determined with ASTRA software.

### Melting curves

SrfE_CTD_, SrfE_del_, and SrfE_4Ala_ in original buffer (10mM Tris HCl pH 8.3, 250mM NaCl, 1mM TCEP, 2% v/v glycerol) were diluted to 1.5 mg/mL, spun down at 14.1K x g for 15 min at 4°C, loaded into capillaries (Prometheus PR-C002) in technical triplicates, and run on a melting ramp from 25-95°C at a rate of 1°C/min on a Prometheus Panta Nanotemper. Traces, derivatives, and melting points were determined with PR.Panta Analysis software version 1.

### Statistics

Statistical analysis was performed using Prism 11 (GraphPad Software). Statistical parameters and significance are reported in figure legends.

