## Supplementary Materials for "A bacterial pathogen reveals a new mechanism of micron-scale nucleocytoplasmic transport"

### Supplemental Figures:

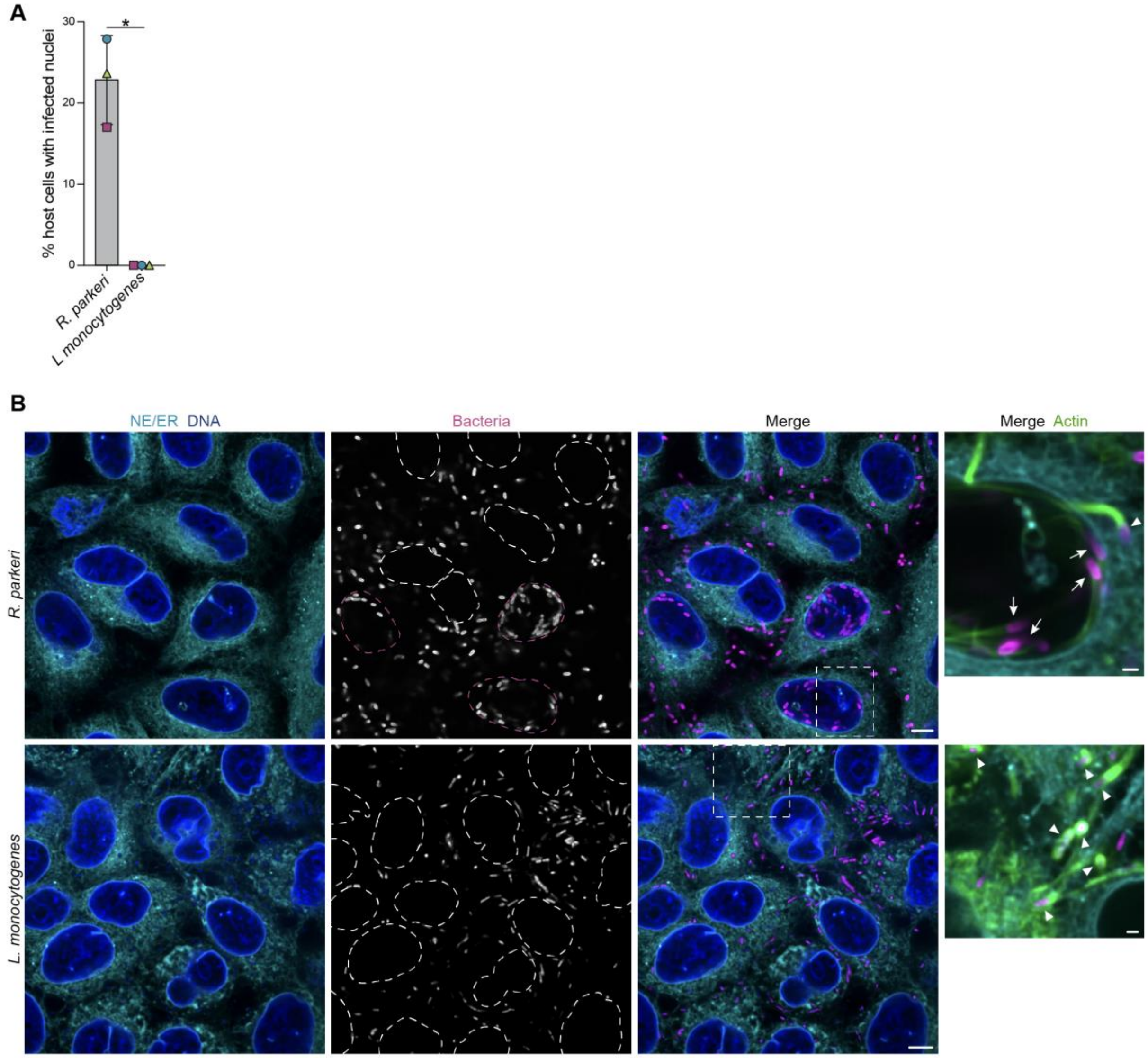


**Fig. S1.**

***Listeria monocytogenes* does not invade human cell nuclei.**

(**A**) Quantification of nuclear invasion in A549 cells infected with *R. parkeri* (72 hpi) or GFP-expressing *L. monocytogenes* (5 hpi). Data are mean ± S.D. from n = 3 biological replicates (>100 infected cells/condition/replicate). Significance calculated via two-tailed Welch's T-test. ns = not significant. (**B**) Representative images from (A), showing host nuclei (DNA in blue, Hoechst), NE/ER (cyan, concanavalin A), indicated bacteria (magenta), and F-actin (green, phalloidin). Nuclear boundaries (dotted outlines), invaded nuclei (dotted magenta outlines). Insets enlarged to right, cytoplasmic bacteria with actin tails (white triangles), nuclear bacteria with actin tails (white arrows) Scale bars, 5 µm. Scale bars of insets, 1 µm. $*P \leq 0.05$.

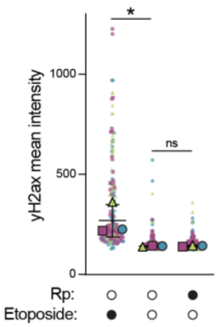


**Fig. S2.**

***R. parkeri* infection does not increase host cell DNA damage response.**

Immunofluorescence detection of DNA damage response (anti-yH2ax) in A549 cells treated with vehicle or 20 μM etoposide for 4 h, or infected with *R. parkeri* (Rp) for ~ 72 h. Mean signal intensity was calculated per cell. Medians from independent experiments (large, outlined shapes) were superimposed over individual mean cell intensities (small shapes). Data are mean ± S.D. from n = 3 biological replicates (>50 infected cells/condition/replicate). Significance calculated via one-way ANOVA multiple comparisons. *$P \leq 0.05$, ns = not significant.

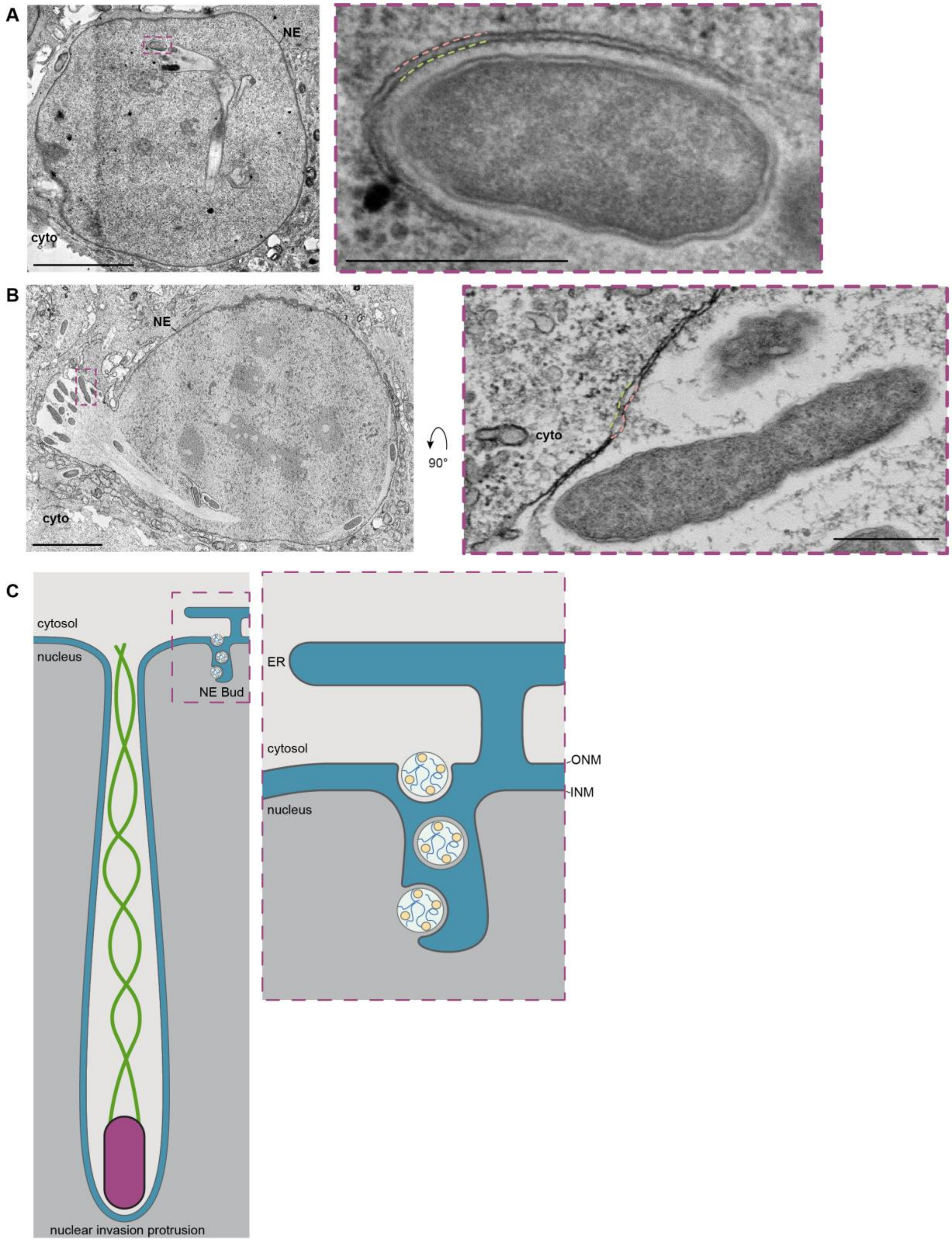
A
NE
cyto
B
NE
cyto
90°
cyto
C
cytosol
nucleus
NE Bud
nuclear invasion protrusion
ER
cytosol
nucleus
ONM
INM

**Fig. S3.**

***R. parkeri* nuclear protrusions are distinct from nuclear envelope buds**

(**A**-**B**) mEmerald-Sec61 U2OS cells infected with BFP-expressing *R. parkeri* for ~72 h were processed for STEM. Representative images showing invading (A) or egressing (B) nuclear protrusions from n = 3 independent experiments, 23 invasion and 6 egress events visualized. Insets (dotted line): enlarged views showing inner (pink) and outer (green) nuclear membranes at protrusions; inset scale bar, 0.5 µm. NE, nuclear envelope; cyto, cytoplasm. Scale bars, 5 µm. (**C**) To-scale comparison of nuclear invasion protrusions used to transport *R. parkeri* (magenta) with an actin tail (green) and nuclear envelope (NE) buds used to transport RNPs (light blue and yellow). NE bud is shown enlarged to the right.

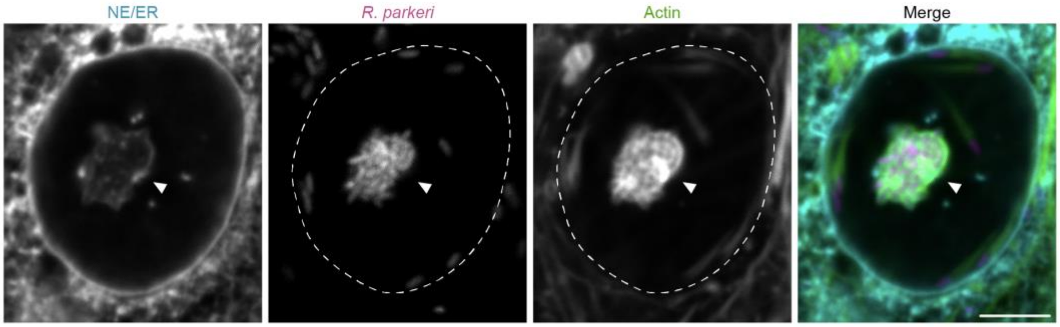


**Fig. S4.**

**Confined actin tails at nuclear invasion protrusions.**
Example image showing bacteria (magenta) with confined actin tails (green) within a nuclear invasion protrusion boundary (cyan) from Fig. 1H. Nuclear boundary (dotted outline), nuclear invasion protrusion (white triangle). Scale bar, 5 µm.

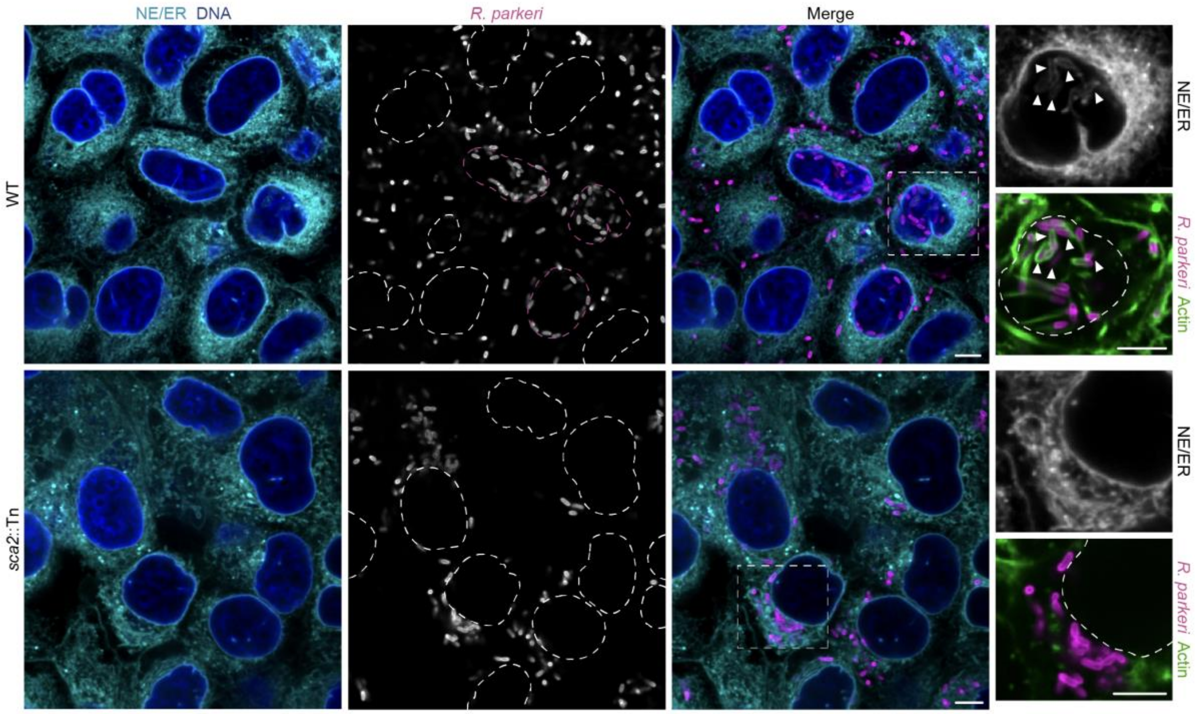


**Fig. S5.**

**Actin-based motility is required for nuclear protrusion formation and invasion.**

Example images from Fig. 1I-J. Nuclear boundaries (dotted outlines), invaded nuclei (dotted magenta outlines). Insets enlarged to right. *R. parkeri* in nuclear invasion protrusions (white triangles). Nuclear boundaries (dotted white outlines). Scale bars, 5 µm.

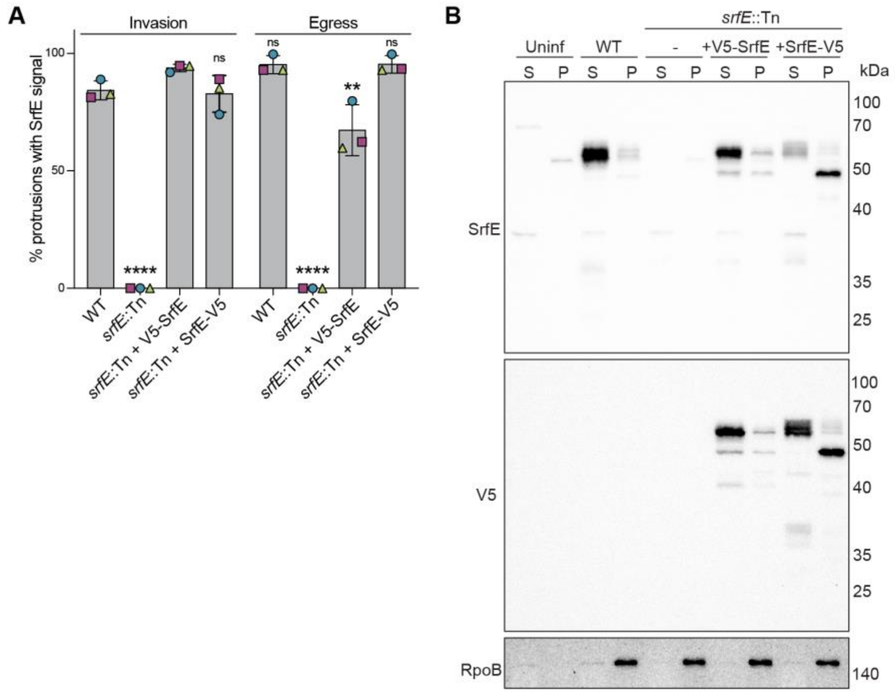


**Fig. S6.**

**SrfE is secreted from WT and SrfE complement *R. parkeri* strains and localizes to nuclear invasion and egress protrusions.**

**(A)** Data from Fig. 2A quantifying SrfE localization to nuclear invasion and egress protrusions in mEmerald-Sec61 U2OS cells infected with WT, *srfE*::Tn, and complement *R. parkeri* strains (BFP, magenta) for ~72 h. Data are mean ± S.D. from n = 3 biological replicates (>50 invasion or >15 egress protrusions/condition/replicate). Significance relative to WT calculated via one-way ANOVA with Dunnett's multiple comparisons test. **(B)** Western blot analysis of lysates from A549 cells infected with indicated *R. parkeri* strains for ~72 h. Cells were selectively lysed to separate supernatants (S), containing secreted effectors, and pellets (P), containing intact *R. parkeri.* SrfE (predicted 48 kDa) was detected with a polyclonal antibody to the endogenous protein or with an anti-V5 antibody. Note the presence of multiple species that are absent from uninfected lanes. DNA-directed RNA polymerase subunit beta (RpoB, 150 kDa) serves as a non-secreted control. Blot is representative of n = 2 biological replicates. ****P ≤ 0.0001, **P ≤ 0.01, ns = not significant.

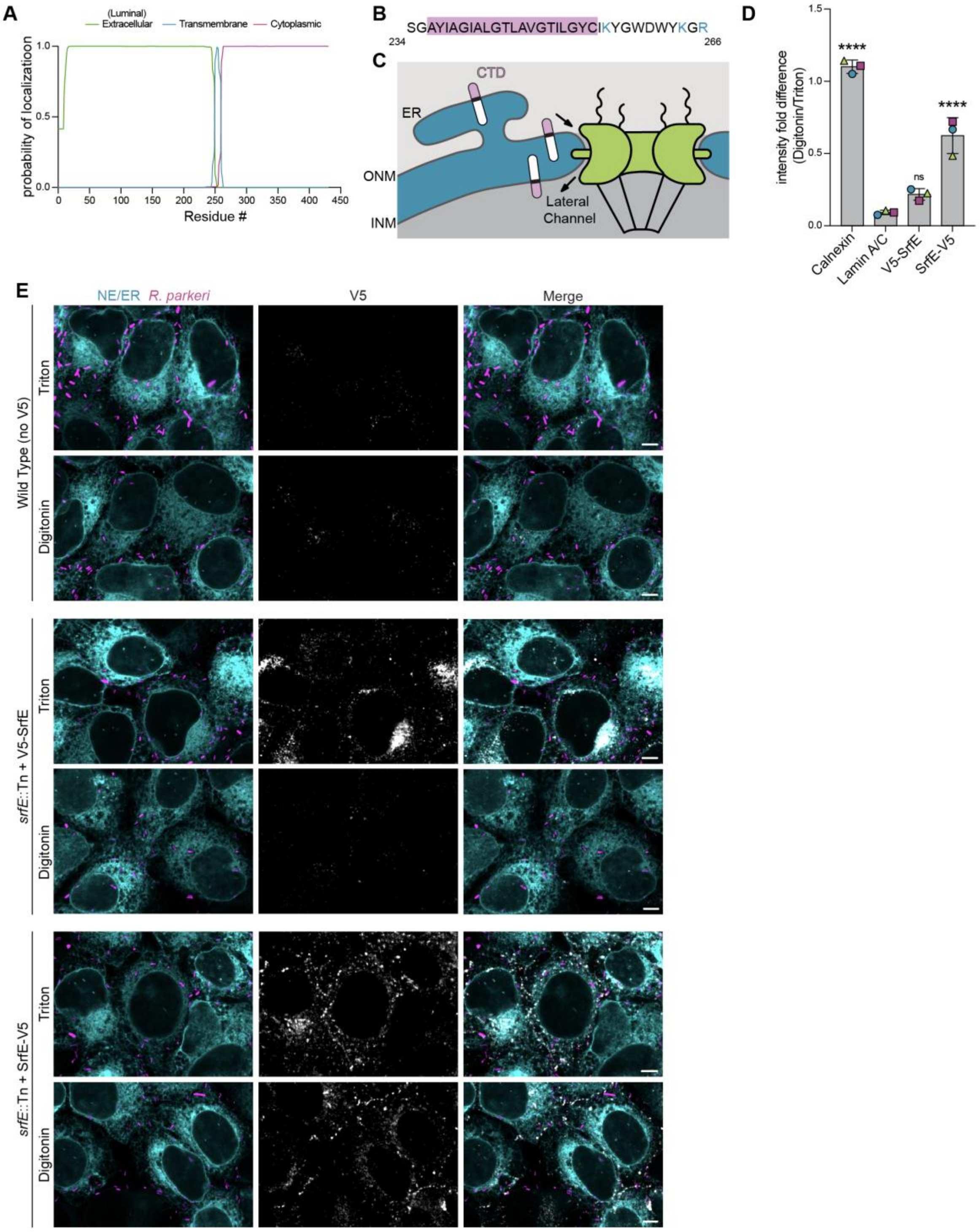


**Fig. S7.**
**SrfE is a Type I transmembrane protein.**
(**A**) PSIPRED (*61*) predicts SrfE contains a single Type I TM domain spanning residues 236-255. (**B**) SrfE residues 234-266; cluster of positively charged residues (cyan) on the C-terminal side of SrfE TM domain (light purple). (**C**) Predicted orientation of SrfE in the endoplasmic reticulum (ER), outer nuclear membrane (ONM), and inner nuclear membrane (INM) based on (A-B) and

experimental results in (D-E). (**D-E**) Quantification (D) and representative images (E) of selective permeabilization assays. mEmerald-Sec61 U2OS cells (cyan) were infected with indicated *R. parkeri* strains (magenta) for ~72 h and stained for V5, lamin A/C, or calnexin accessibility (gray). WT provided background V5 immunofluorescence intensities. Images were analyzed by measuring the mean intensity and subtracting background values. Data are mean ± S.D. for n = 3 biological replicates (>50 infected cells/condition/replicate for V5-SrfE and SrfE-V5 expressing strains and >20 protrusions/condition/replicate for lamin and calnexin controls). Significance, relative to Lamin A/C was calculated via one-way ANOVA with Dunnett's multiple comparisons test. Scale bar, 5 µm. ****$P \leq 0.0001$, ns = not significant.

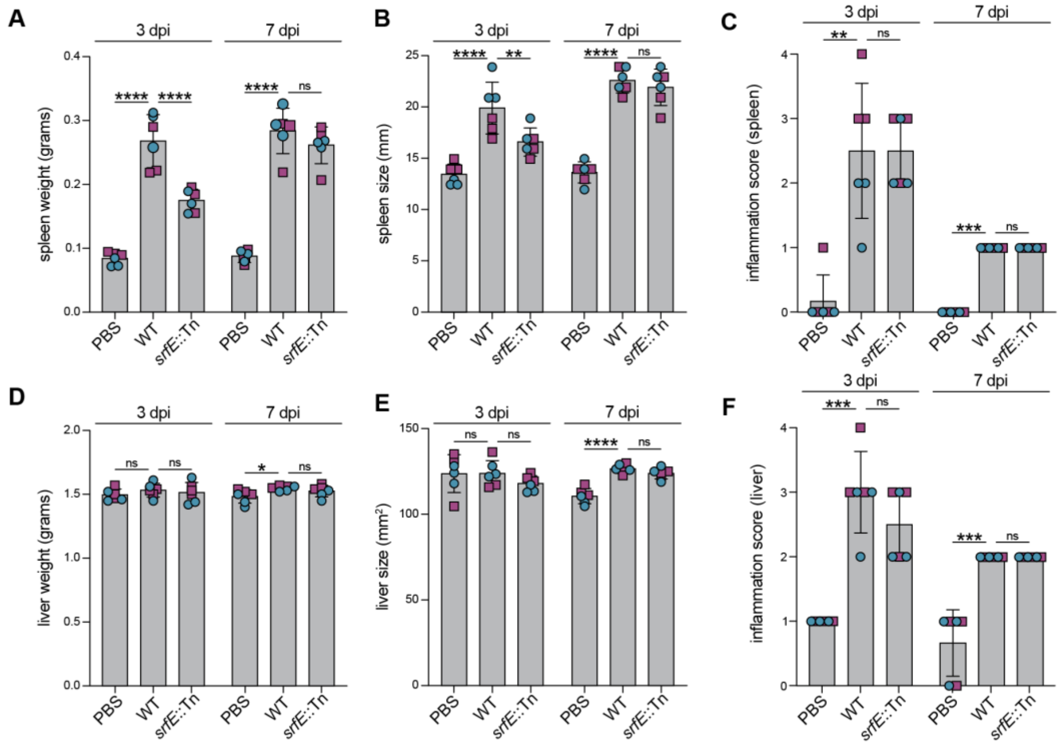


**Fig. S8.**

**The effect of SrfE on mouse organ size, weight, and inflammation during *R. parkeri* infection.** (**A-B, D-E)** Spleen weight (A), spleen size (B), liver weight (D), and liver size (E) from mice infected with the indicated *R. parkeri* strains for 3 and 7 days. Organs were collected and measured at indicated time points. Data are mean ± S.D. from n = 2 biological replicates (3 mice/condition/replicate). Significance calculated via one-way ANOVA with Dunnett's multiple comparisons test, relative to WT. (**C, F**) Spleen (C) and liver (F) inflammation from mice infected with the indicated *R. parkeri* strains for 3 and 7 days. Organs were collected and processed for staining with hematoxylin and eosin (H&E) in 5 µm-thick sections. Sections were scored semi-quantitatively for inflammation by a board-certified veterinary pathologist on a 5 point ordinal scale: 0 = absent, 1 = rare inflammatory cells (minimal), 2 = involvement of <25% of the tissue (mild), 3 = involvement of 25-50% of the tissue (moderate), and 4 = involvement of >50% of the tissue (marked). Data are mean ± S.D. from n = 2 biological replicate (3 mice/condition/replicate). Significance calculated via one-way ANOVA with Kruskal-Wallis multiple comparisons test, relative to WT. ns = not significant. ****P ≤ 0.0001, ***P ≤ 0.001, **P ≤ 0.01, *P ≤ 0.05, ns = not significant.

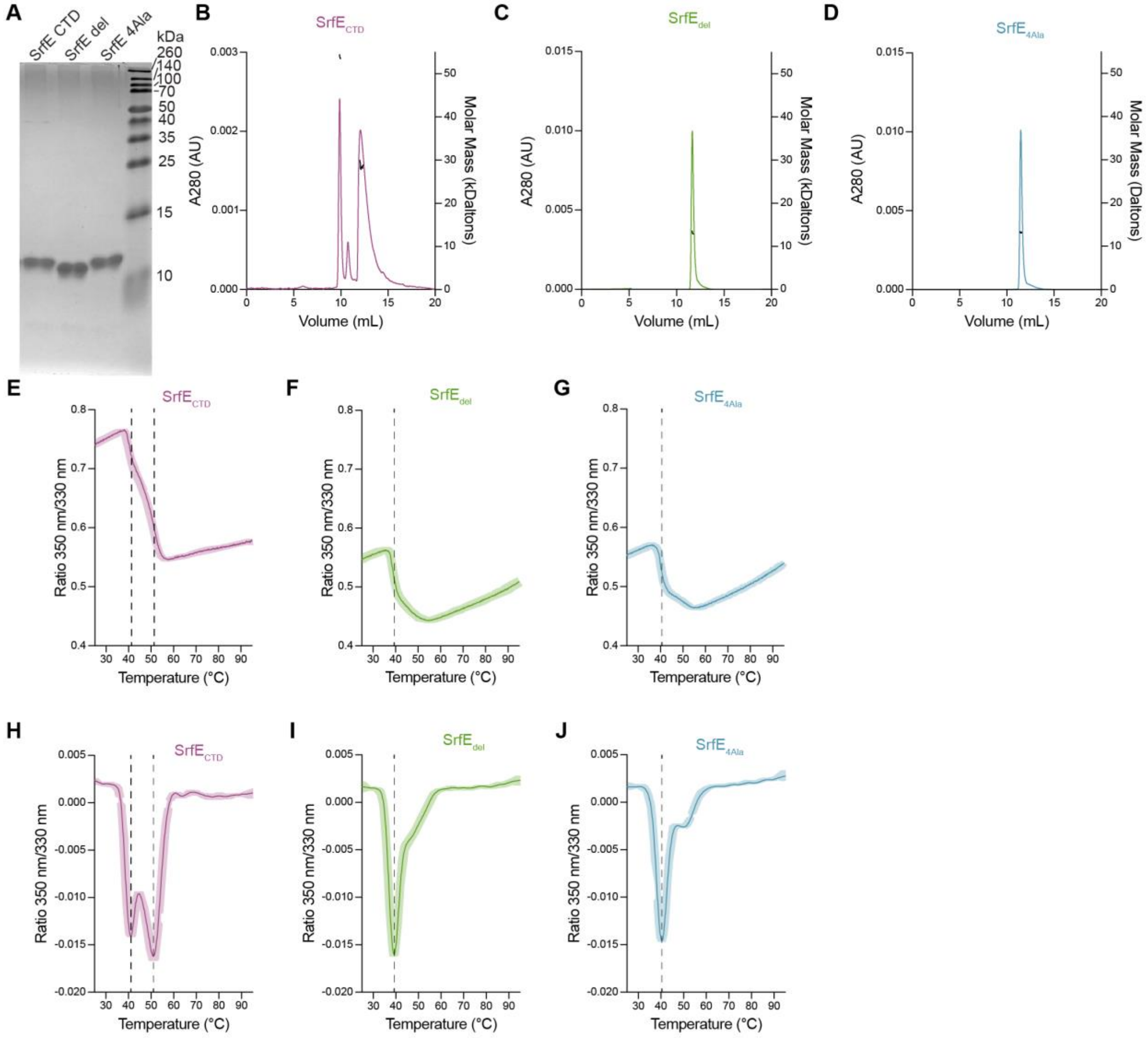


**Fig. S9.**

**$\mathrm{SrfE_{CTD}}$ is a tetramer *in vitro***

(**A**) Purified protein constructs used for assays on a Coomassie gel. Constructs were diluted to 25 μM and 2.5 μL was run in each lane. Gel is representative of n = 2 biological replicates. (**B**-**D**): Elution profiles (colored traces) and molar masses (black traces) of purified proteins as determined by size exclusion chromatography multi angle light scattering (SEC-MALs). Purified proteins at 0.5 mg/mL were injected onto a WTC-030S5 size exclusion chromatography analytical column. Molar masses were calculated with ASTRA software. Graphs representative of n = 3 biological replicates. (**E**-**J**): Melting curves of purified proteins measured as ratio of natural absorbance at 350 and 330 nm (E-G) or a derivative of these values (H-J) over a temperature gradient. A melting ramp of 1°C/min from 25-95°C was run on purified protein at 1.5 mg/mL on a Prometheus Panta nanotemper. Data are mean (lines) ± S.D. (shaded area) from n = 3 technical replicates. Graphs representative of n = 3 biological replicates. Vertical dotted lines indicate inflection points indicative of an unfolding event calculated by PR.Panta Analysis version 1.

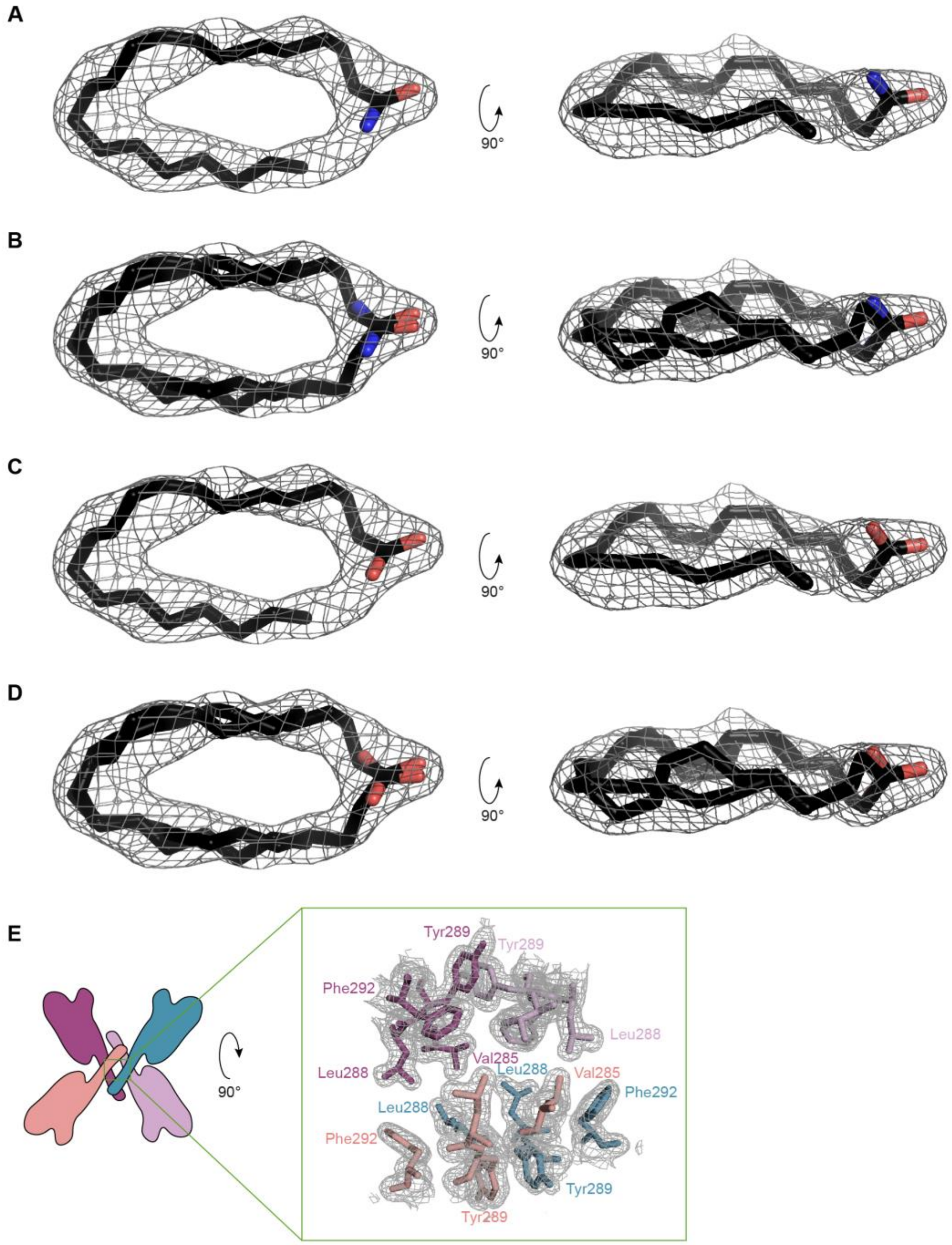


**Fig. S10.**
**Fatty acid binding and tetramerization interface of SrfE$_{CTD}$**
(**A-D**) One (A, C) or two copies (B, D) of arachidonoyl-2-fluoroethylamide with the 2-fluoro-ethyl group removed (A-B) or arachidonic acid (C-D) docked into electron density (light gray) from top and side angle. Carbon in black, oxygen in red, nitrogen in blue. (**E**) Tetramerization interaction interface. Residues involved in interaction shown as sticks, colored as in Fig. 4A, docked into electron density (light gray mesh). Not all residues labeled due to space.

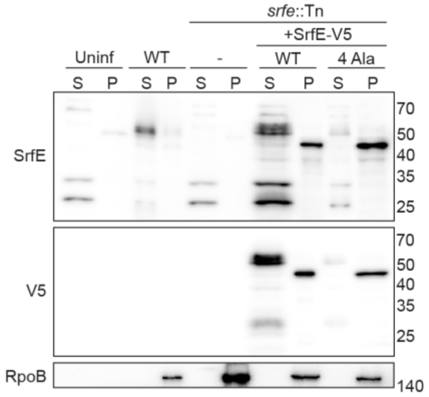


**Fig. S11.**
**Mutating the tetramerization interface of SrfE leads to impaired secretion.**
Western blot analysis of lysates from A549 cells infected with indicated *R. parkeri* strains for ~72 h. Cells were selectively lysed to separate supernatants (S), containing secreted effectors, and pellets (P), containing intact *R. parkeri.* SrfE (predicted 48 kDa) was detected with a polyclonal antibody to the endogenous protein or with an anti-V5 antibody. Note the presence of multiple species that are absent from uninfected lanes. DNA-directed RNA polymerase subunit beta (RpoB, 150 kDa) serves as a non-secreted control. Blot is representative of n = 2 biological replicates.

**Table S1.**

| **Condition** | **dpi** | **Liver** | **Spleen** |
|---|---|---|---|
| PBS | 3 | 0/6 | 0/6 |
| WT | 3 | 5/6 | 2/6 |
| *srfE*::Tn | 3 | 2/6 | 1/6 |
| PBS | 7 | 0/6 | 0/6 |
| WT | 7 | 1/6 | 0/6 |
| *srfE*::Tn | 7 | 0/6 | 0/6 |

Fraction of samples with thrombosis/total for each tissue at the indicated days post infection (dpi).

**Table S2.**

| **PDB Accession Code** | **35YA** |
|---|---|
| **Data Collection** | |
| Space group | *P1* |
| Unit cell parameters (Å; º) | $a = 45.45,\ b = 126.26,\ c = 126.76;$ |
| | $\alpha = 119.67,\ \beta = 90.05,\ \gamma = 90.21$ |
| Resolution range (Å) | 30.00 - 1.85 (1.88 - 1.85) |
| No. of reflections | 204,650 (9,496) |
| $R_{merge}$ (%) | 6.9 (50.3) |
| $CC_{1/2}$ | 0.995 (0.424) |
| Completeness (%) | 97.3 (90.2) |
| $\langle I/\sigma(I)\rangle$ | 15.5 (1.8) |
| Multiplicity | 3.5 (2.9) |
| Wilson *B* factor | 24.8 |
| **Refinement** | |
| Resolution range (Å) | 28.60 - 1.85 (1.88 - 1.85) |
| Completeness (%) | 97.2 (90.0) |
| No. of reflections | 193,396 (9,423) |
| $R_{work}/R_{free}$, (%) | 16.1/18.8 (22.2/26.3) |
| Protein chains/atoms | 12/12426 |
| Ligand/Solvent atoms | 294/2,001 |
| Mean temperature factor (Å$^2$) | 29.0 |
| **Coordinate Deviations** | |
| R.m.s.d. bonds (Å) | 0.004 |
| R.m.s.d. angles (º) | 1.345 |
| **Ramachandran plot** | |
| Favored (%) | 99.0 |
| Allowed (%) | 1.0 |
| Outside allowed (%) | 0.0 |

Refinement statistics for structure of $SrfE_{CTD}$. Values in parentheses are for the outer shell. Completeness values before twin corrections were applied.

**Table S3.**

| Compounds | LogFC | Mass | Molecule | Lipid(Y/N) |
|---|---|---|---|---|
| (+)-Prosopinine Esi+8.991 | 4 | 335.2432 | $C_{16}H_{33}NO_3$ | Y |
| *N*-(2-fluro-ethyl) arachidonoyl amine Esi+9.185 | 4 | 335.2439 | $C_{22}H_{36}FNO$ | Y |
| *N*-(2-fluro-ethyl) arachidonoyl amine Esi+8.922 | 4 | 331.2711 | $C_{22}H_{36}FNO$ | Y |
| (+)-Prosopinine Esi+8.763 | 4 | 331.2717 | $C_{16}H_{33}NO_3$ | Y |
| (+)-Prosopinine Esi+8.999 | 4 | 335.2432 | $C_{16}H_{33}NO_3$ | Y |
| (+)-Prosopinine Esi+9.61 | 4 | 295.2512 | $C_{16}H_{33}NO_3$ | Y |
| C18 H35 N O4 Esi+8.998 | 4 | 295.2509 | $C_{18}H_{35}NO_4$ | Y |
| (4OH, 8Z, t18:1) sphingosine Esi+10.16 | 4 | 329.2568 | $C_{18}H_{37}NO_2$ | Y |
| 13,14-dihydroxy-docosanoic acid Esi+10.614 | 4 | 297.2668 | $C_{22}H_{44}O_4$ | Y |
| (+)-Prosopinine Esi+9.041 | 3.420956 | 371.3144 | $C_{16}H_{33}NO_3$ | Y |

Top 10 compounds identified through mass spectrometry analysis of protein crystals of $SrfE_{CTD}$. For all data see Raw data file 1. LogFC=LogFoldChange abundance compared to crystallization buffer.

**Movie S1.** View through and modeled representation of FIB-SEM volume of an *R. parkeri*-infected A549 cell nucleus. Bacteria, magenta; nuclear envelope, cyan (related to Fig. 1C).

**Movie S2.** *R. parkeri* (magenta) movement within the nucleus of a U2OS mEmerald-Sec61beta cell (nuclear envelope and endoplasmic reticulum in cyan) (related to Fig. 1). Data were obtained as in Fig 1. E-F. Time, hours:min.

**Movie S3.** Multiple *R. parkeri* bacteria (magenta) invade the nucleus of a U2OS mEmerald-Sec61beta cell (nuclear envelope and endoplasmic reticulum in cyan) (related to Fig. 1E). Bacteria begin in a single, large protrusion, and enter the nucleoplasm by the end of the movie. Data were obtained as in Fig 1. E-F. Time, hours:min.

**Movie S4.** Multiple *R. parkeri* bacteria (magenta) exit the nucleus of a U2OS mEmerald-Sec61beta cell (nuclear envelope and endoplasmic reticulum in cyan) (related to Fig. 1F). Bacteria begin in multiple egress protrusions and indicated groups enter the cytoplasm by the end of the movie. Data were obtained as in Fig 1. E-F. Time, hours:min.

**Movie S5.** Individual *R. parkeri* bacteria (magenta) invade and exit the nucleus of a U2OS mEmerald-Sec61beta cell (nuclear envelope and endoplasmic reticulum in cyan) (related to Fig. 1E-F). Cell begins and ends with no nuclear envelope protrusions. Data were obtained as in Fig 1. E-F. Time, hours:min.
